# PRMT5 activity sustains histone production to maintain genome integrity

**DOI:** 10.1101/2025.07.03.663002

**Authors:** Jacob S. Roth, Joseph D. DeAngelo, Dejauwne L. Young, Maxim I. Maron, Ankita Saha, Hugo Pinto, Varun Gupta, Noah Jacobs, Subray Hegde, Jennifer T. Aguilan, Joel Basken, Joey Azofeifa, Charles C. Query, Simone Sidoli, Arthur I. Skoultchi, David Shechter

## Abstract

Histone proteins package DNA into nucleosomes, forming chromatin and thereby safeguarding genome integrity. Proper histone expression is essential for cell proliferation and chromatin organization, yet the upstream regulators of histone supply remain incompletely understood. PRMT5—a cell essential type II protein arginine methyltransferase frequently overexpressed in cancer—catalyzes symmetric dimethylation of arginine residues. Using time-resolved nascent transcriptional profiling, quantitative proteomics, and imaging, we show that PRMT5 activity is required to sustain histone transcription and histone protein synthesis during S phase. PRMT5 inhibition or knockdown leads to rapid histone mRNA depletion, loss of histone proteins, and accumulation of replication-associated nuclear abnormalities. We further show that soluble histone H4 accumulates at histone locus bodies (HLBs) upon PRMT5 inhibition, and that PRMT5-substrate H4 Arginine 3 mutants localize more robustly to HLBs than do wildtype H4. These findings support a model in which PRMT5-mediated methylation of histone H4 regulates histone transcription. Our findings establish PRMT5 as a central coordinator of histone homeostasis and provide a mechanistic rationale for its essential role in proliferating cells.

## Introduction

In eukaryotes, DNA is packaged into chromatin, a dynamic superstructure that both protects the genome and regulates its accessibility to transcription, replication, and repair machinery. Chromatin is composed of repeating nucleosomes, each consisting of 146 base pairs of DNA wrapped around a hetero-octamer of the four core histones H2A, H2B, H3, and H4^5^. The linker histone H1 binds to DNA between nucleosomes, promoting chromatin compaction and regulating higher-order structure by controlling nucleosome spacing^6, 7^.

During each cell cycle, proliferating cells duplicate their genome and produce a corresponding stoichiometric supply of new histones to preserve genomic stability and maintain chromatin organization^8^. Insufficient histone synthesis disrupts nucleosome assembly and can lead to replication stress, DNA damage, transcriptional dysregulation, and cell death^9-12^. To prevent such outcomes, cells couple histone biosynthesis to DNA replication through tightly regulated transcriptional and posttranscriptional control mechanisms^13-15^.

Replication-dependent histone genes are encoded in multiple copies and organized into two main clusters: 55 genes in the chromosome 6 HIST1 cluster, and six genes in the chromosome 1 HIST2 cluster^16^. These gene clusters are regulated through the formation of histone locus bodies (HLBs), membraneless nuclear compartments enriched in histone gene transcription and processing machinery^17^. Upon entry into S-phase, Cyclin E/CDK2-dependent phosphorylation of NPAT stimulates transcription of histone genes^17-20^. Histone pre-mRNAs are transcribed without 3’ polyadenylation and undergo specialized 3′ end processing in the HLB prior to cytoplasmic export and translation^21^. Histone chaperones then escort nascent histone proteins into the nucleus and participate in their deposition into chromatin^22, 23^.

Protein arginine methyltransferase 5 (PRMT5) catalyzes the symmetric dimethylation (Rme2s) of arginine residues on hundreds of substrates, including spliceosomal proteins, chromatin remodelers, and transcriptional regulators^24, 25^. PRMT5 is essential for development, stem cell maintenance, and cell cycle progression^26-29^. Its expression is frequently elevated in cancers—including glioblastoma, lymphoma, lung, and breast cancers—and correlates with poor prognosis and aggressive disease^30, 31^. These observations motivated the development of selective PRMT5 inhibitors, several of which are currently in clinical trials for both solid and hematologic malignancies^32, 33^. Due to its many substrates and numerous biological functions, a unifying explanation for PRMT5’s essential role in proliferating cells remains elusive. While PRMT5 has been implicated in regulating pre-mRNA splicing, DNA damage repair, and transcriptional repression^34-39^, none of these functions explain the vertebrate cell essential nature of PRMT5 activity.

To understand the cellular roles of PRMT5 activity, here we dissect the immediate consequences of PRMT5 inhibition. Using an integrated approach combining nascent transcriptional profiling, quantitative and time-resolved proteomics, and high-resolution imaging, we uncover an unexpected role for PRMT5 in directly sustaining S-phase histone gene transcription and concomitant maintenance of genome integrity. We further test the histone transcription feedback regulatory role of PRMT5 and its substrate histone H4 at the HLB. Our study establishes PRMT5 as a critical regulator of histone homeostasis and genome stability, offering a unifying explanation for its essentiality in proliferating cells and rationale for its inhibition in cancer therapy.

## Results

### PRMT5 disruption induces γH2AX containing micronuclei

To determine the critical cellular pathways that are dependent on PRMT5 activity and to identify direct consequences of inhibition, we treated p53-wildtype A549 lung adenocarcinoma cells with selective PRMT5 inhibitors and explored cellular consequences over multiple days of treatment. In these studies, we used both peptide-competitive GSK591 (also known as GSK3203591 and EPZ015866) and SAM-competitive (LLY-283) PRMT5 inhibitors along with their matched negative controls (SGC2096 and LLY-284, respectively) (**Supplementary Figure 1a**). With two independent modes of inhibition, this toolbox allowed precise and robust, rapid timescale investigation of PRMT5 cellular activity.

Immunoblot and RT-qPCR analyses revealed that p21/CDKN1A, a downstream target of p53, was induced after 12-24 hours of PRMT5 inhibition, considerably earlier than previously reported^36^(**Figure 1a and Supplementary Figure S1b**). This was coincident with an increase in the DNA damage marker γH2AX, loss of total symmetric dimethylarginine, chromatin accumulation of the PRMT5 substrate SNRPB, and an increase in H3K27me3 (**Figure 1a-b and Supplementary Figure S1c**). As prior studies showed that PRMT5 knock-down activated p53 indirectly via alternative splicing of its negative regulator MDM4^36^, we tested the timing of MDM4 differential splicing after PRMT5 chemical inhibition. Both intron-spanning RT-PCR and qPCR quantification revealed that alternative splicing of MDM4 was detectable within hours of PRMT5 inhibition by GSK591 (**Supplementary Figure S1d,e**). This effect was reproduced by the SAM-competitive inhibitor LLY-283, but not by structurally similar negative control compounds LLY-284 or SGC2096, confirming that rapid MDM4 splicing is a direct consequence of PRMT5 catalytic inhibition.

**Figure 1.**
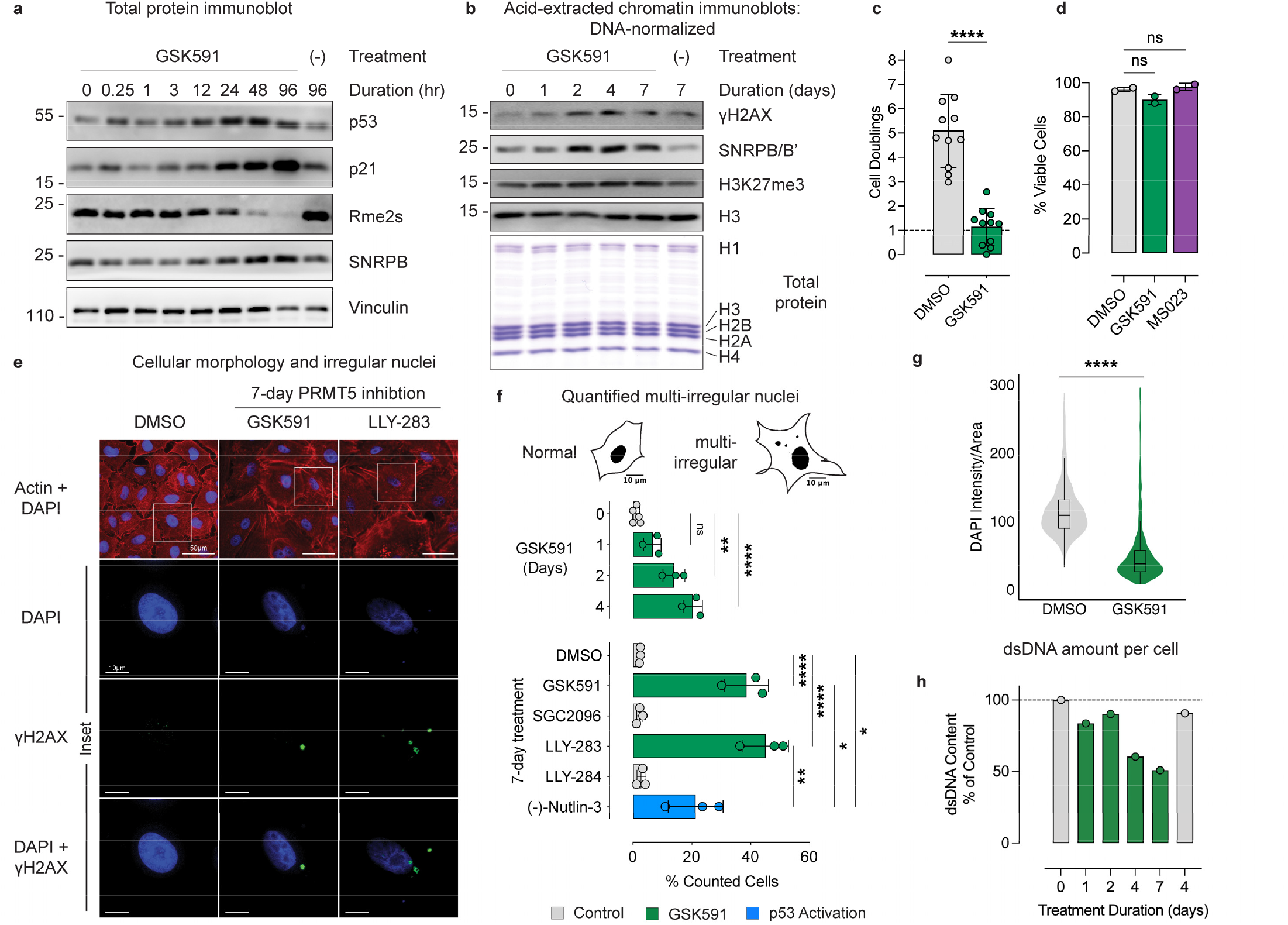
PRMT5 disruption induces γH2AX containing micronuclei. **a)** Immunoblots of total cellular lysate after PRMT5 inhibition. **b)** Immunoblots of acid-extracted chromatin after PRMT5 inhibition, loading normalized by DNA content. **c)** Cell count of A549 cells after 4-day PRMT5 inhibition. **d)** Trypan blue viability of A549 cells after 4-day PRMT5 inhibition. **e)** A549 cells treated with 7-days of PRMT5 inhibition, highlighting enlarged “fried-egg” appearance of DAPI-stained nuclei (blue) in actin (red). Immunofluorescence staining of γH2AX (green) overlaid upon nuclei (blue), highlighting γH2AX-positive micronuclei separate from the primary nucleus. Scale bar 50μm in Actin+DAPI and 10μm in inset images. **f)** Quantified multi-irregular nuclei observed after treating A549 cells with 0 to 4 days of PRMT5 inhibition and 7 days of PRMT5 inhibition, PRMT5-inhibitor negative controls, or p53 activation by (-)-Nutlin3a. Inset images of representative “Normal” (7-day DMSO) and “Multi-irregular” (7-day GSK591) nuclei; DAPI thresholding in FIJI to highlight nuclei with actin outline. **g)** Ratio of DAPI intensity per nucleus area (RFU/μm^2^) from 7-day PRMT5-inhibited cells; >100 cells were analyzed for each of three biological replicates in each treatment. **h)** TRIzol-extracted and quantified dsDNA from equivalent cell number; n=1 biological replicate. Student’s one-tailed t-test was applied; p-value = *<0.05, **<0.005, ***<0.0005, ****<0.00005, n.s. = not significant.

Together, these results demonstrated that PRMT5 inhibition rapidly disrupts p53 regulatory pathways, well before overt phenotypic changes become apparent. Although p53 activation was evident within 24 hours, cells did not exhibit reduced metabolic activity by MTS assay until 72 hours following PRMT5 inhibition (**Supplementary Figure S1f**). However, direct cell counting revealed a more immediate and profound proliferative defect: while control cells underwent an average of 5.1 doublings over four days, PRMT5-inhibited cells completed only a single division (**Figure 1c**). Despite this marked growth arrest, there was no evidence of cell death: GSK591-treated cells did not die nor did control cells following treatment with the Type I PRMT inhibitor MS023, which inhibits asymmetric dimethylation (Rme2a) of arginine (**Figure 1d**). We concluded that that PRMT5 inhibition induces a non-lethal restriction on proliferation.

Our prior work reported that PRMT5 inhibition in A549 cells induced an enlarged, “fried egg” phenotype, with increases both to cellular size and the number of nuclei per cell^35^. This phenotype was consistent across multiple modes of PRMT5 disruption, including both peptideand SAM-competitive inhibition by GSK591 and LLY-283, respectively, and also by CRISPRi-mediated PRMT5 genetic knockdown^40^(**Figure 1e and Supplementary Figure S1g**). We also noted a similar response in the non-cancerous BEAS-2B cell line (*not shown*). GSK591-treated cells exhibited a dose and time-dependent increase in β-galactosidase staining, which, together with their enlarged size and slow growth is consistent with cellular senescence (**Supplementary Figure S1h,i**).

To assess whether these cellular effects were direct consequences of PRMT5 inhibition or indirect via p53 activation, we compared the morphology of cells treated with the p53-activator (-)-Nutlin-3 to those treated with PRMT5 inhibitors. After seven days of treatment, both conditions produced similarly enlarged cells. However, we observed a striking difference in nuclear morphology: PRMT5-inhibited cells frequently displayed multiple, irregular, and fractured nuclei, a feature greatly depleted in (-)-Nutlin-3-treated cells (**Supplementary Figure S1g**).

The irregular nuclear structures in PRMT5-inhibited cells included micronuclei, a hallmark of mitotic catastrophe^41^. Micronuclei typically exhibit double stranded DNA breaks and subsequent phosphorylation of S139 on histone H2AX (γH2AX). PRMT5-inhibited cells exhibited an increase in γH2AX on acid extracted histones after two days of treatment (**Figure 1b**). After probing these cells by γH2AX immunofluorescence (IF), we found that the signal was restricted to discrete micronuclear foci. A profile plot line for a representative micronucleus from 7-days of GSK591 highlights the overlapping nuclear and γH2AX signals (**Supplementary Figure S1j**). This observation was in stark contrast to the intense and diffuse nuclear staining found after doxorubicin treatment (**Supplementary Figure S1k**).

We quantified nuclei from treated cells, annotating them as: 1) mononuclear; 2) binucleate (two equal-sized nuclei); or 3) multi-irregular (containing micronuclei, nuclear buds, or more than one nucleus of unequal size). The most prominent type of multi-irregular nuclei in PRMT5-inhibited cells contained micronuclei. PRMT5-inhibited cells accumulated multi-irregular nuclei over time, containing ∼8x more multi-irregular nuclei (40%) than control cells after seven days of treatment (**Figure 1f**). Neither SGC2096 nor LLY-284 negative control compounds differed from DMSO-treated control. p53 activation via (-)-Nutlin-3 was confirmed to induce p21 expression but only increased multi-irregular nuclei to 20% of observed cells, suggesting that p53 activation alone does not explain the impact of PRMT5-inhibition on cell nuclei (**Supplementary Figure S1l and Figure 1f**). An increase in multi-irregular cells was observed for PRMT5 knockdown in A549 cells and 4-day PRMT5 inhibition in non-cancerous lung epithelial BEAS-2B cells (**Supplementary Figure S1m**).

As we observed a stark reduction in DAPI intensity in all cases of PRMT5 disruption, we quantified DAPI intensity levels in images from 4-day GSK591-treated cells and found a significant decrease (**Figure 1g**). To test if total cellular DNA content was reduced after PRMT5 inhibition, cell-count normalized samples were extracted with TRIzol and dsDNA was quantified by Qubit assay. This revealed a time-dependent loss of DNA content over the course of 7 days of PRMT5 inhibition (**Figure 1h**). Together, these results demonstrate that PRMT5 activity is essential for maintaining genome integrity, and its disruption leads to nuclear abnormalities and DNA loss.

### Nascent transcriptomics identifies a PRMT5 activity requirement for histone transcription

Micronuclei can arise in response to diverse cellular stresses, prompting us to investigate which pathways were disrupted by PRMT5 inhibition to drive their formation. To profile initial cellular responses that might precede downstream phenotypes and determine if loss of PRMT5 activity caused changes in gene expression earlier than we and others previously tested, we performed nascent RNA analysis with Precision Run On Sequencing (PRO-Seq). PRO-seq quantifies nascent transcription with base-pair resolution of RNA Polymerase II activity and can observe all transcribed RNAs. After treating cells for 15 minutes, 3 hours, or 2-, 4-, and 7-days of PRMT5 inhibition, we probed their immediate transcriptional status (**Supplementary Figure S2a, Supplementary Table S1**). This analysis revealed a cascade of transcriptomic changes, with discrete sets of genes being altered at 2-, 4-, and 7-days (**Figure 2a**). Principal component analysis of the PRO-seq proteincoding and lncRNA gene-body transcriptome revealed a temporal dependence in PC1-variance over PRMT5 inhibition (**Figure 2b**). Mirroring our prior observations that major cellular phenotypic changes do not arise until after two days of inhibition, the transcriptomes of cells treated for greater than two days of inhibition clustered with pronounced PC1 variance from the control. The main PC2 variance was observed in changes over 7-day PRMT5 inhibition. While the 15 min timepoint clustered with the control condition, the three-hour inhibition condition variance clustered away from control in both PC1 and PC2, suggesting rapid transcriptional changes upon PRMT5 inhibition.

**Figure 2.**
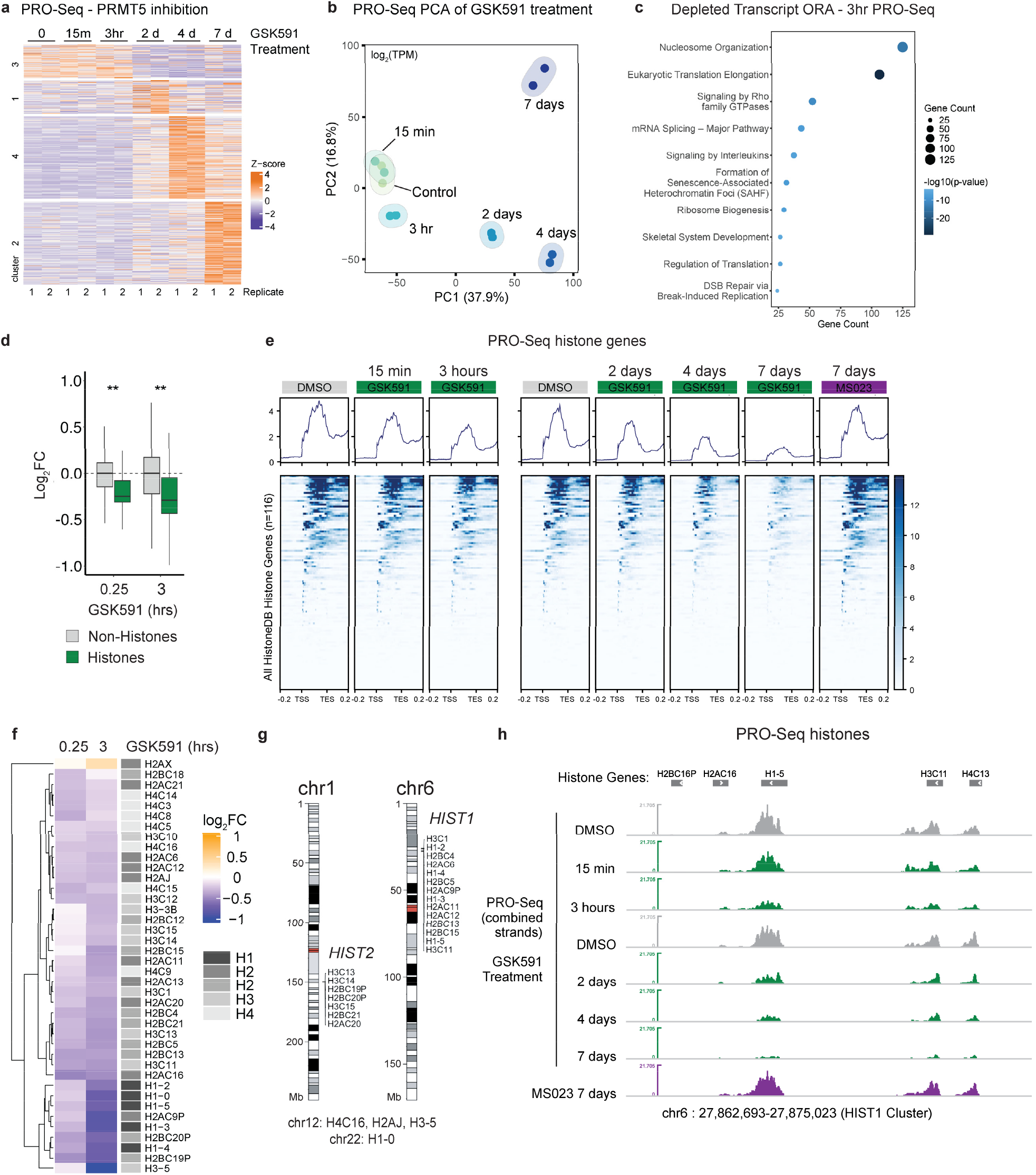
Nascent transcriptomics identifies PRMT5 activity is required for histone transcription. **a)** Heatmap comparing changes in nascent transcription over the duration of PRMT5 inhibition. Color corresponds to Z-score corrected transcripts per million (TPM). **b)** PCA analysis of PRO-seq data. **c)** Over enrichment analysis of transcripts reduced in transcription after 3-hours of PRMT5 inhibition. Enrichment completed with Metascape^2^. **d)** Random permutation testing of Log_2_FC nascent transcription response for all histone genes vs all other genes at 15 minutes and 3-hours of PRMT5 inhibition (100,000 permutations). p-value = **<0.005. **e)** Metaplot of average PRO-seq signal across all histone gene bodies −0.2 kilobases preceding the transcription start site and +0.2 kilobases following the transcription end site (TES). Histone genes retrieved from HistoneDB; plots made with deep-Tools. **f)** Heatmap plotting the log_2_FC of all histone genes observed at 15 min and 3-hours of PRMT5 inhibition via PRO-seq. **g)** Schematic of the replication-dependent histone genes with reduced transcription at 3-hours after PRMT5-inhibition, highlighting the histone clusters HIST1 and HIST2 on chromosomes 6 and 1, respectively. Plots made with KaryoploteR^4^. **h)** Read density (combined PRO-seq strands) at representative histone genes within the HIST1 cluster on chromosome 6.

As a detectable loss of symmetric dimethylarginine (Rme2s) by immunoblot required ≥24 hours of PRMT5 inhibition, the immediacy of these transcriptomic changes was unexpected. We therefore hypothesized that these rapid PRMT5-activity dependent changes might drive the downstream cellular responses we observe upon PRMT5 inhibition. To test this hypothesis and identify transcripts dependent on PRMT5 activity, we performed over-enrichment analysis on transcripts depleted in PRO-seq after three hours of PRMT5 inhibition; this revealed an enrichment for transcripts coding for proteins involved in nucleosome organization (**Figure 2c**). The enriched cluster was populated with genes from a broad set of histone genes, including all core and linker histones and excluding the DNA damage–associated variant H2AX. We tested the PRO-Seq data of all histone genes present in the dataset (n = 40, as defined by histoneDB 2.0^42^) vs. all non-histone genes. Random permutation testing confirmed that the collective downregulation of histone transcripts was significant, both at three hours and as early as 15 minutes following PRMT5 inhibition (**Figure 2d and Supplementary Figure S2b**). RT-qPCR testing confirmed a reduction in bulk random-hexamer primed RNA levels for the two representative linker histones H1-3 and H1-5 after only 24-hours of PRMT5 inhibition (**Supplementary Figure S2c**). Average PRO-seq density was lost across the entire gene body of all histones as early as 15-min following PRMT5 inhibition and persisted throughout the 7 days of the study (**Figure 2e**). Importantly, Type I PRMT inhibition by MS023 did not impact nascent histone transcription (**Figure 2e**).

The histone genes altered after three hours of PRMT5 inhibition are found within the HIST1 and HIST2 clusters on chromosomes 6 and 1, respectively (**Figure 2f,g**). The rapid loss of transcriptional activity at histone genes following PRMT5 inhibition is shown for selected histones within the HIST1 cluster, including H1-5, H3C11, and H4C13 (**Figure 2h**). We conclude that PRMT5 activity contributes to nascent histone gene transcription.

### PRMT5 inhibition phenocopies histone transcription arrest caused by HINFP knockdown

Given that histone gene transcription is immediately reduced upon PRMT5 inhibition, we hypothesized that PRMT5 activity may directly support histone gene expression. We further hypothesized that the transcriptomic and phenotypic cellular responses we observed at two days of PRMT5 inhibition and beyond could be indirect responses to the initial cellular consequences of PRMT5 inhibition, potentially a direct consequence on histone transcription. To investigate how PRMT5 regulates histone transcription, we performed transcription factor enrichment analysis (TFEA^1^) on our PRO-seq dataset (**Supplementary Table S2**). This analysis revealed an enrichment between transcripts downregulated by 3 hours of PRMT5 inhibition and those regulated by HINFP, a transcription factor shown to primarily control expression of histone H4 and select H1 genes (**Figure 3a,b**). To characterize the regulated genes, we plotted publicly available HINFP and NPAT ChIP-seq binding profiles from U2OS cells with the PRO-seq dataset after PRMT5 inhibition, confirming binding of HINFP and NPAT to multiple histone genes (**Figure 3b**).

**Figure 3.**
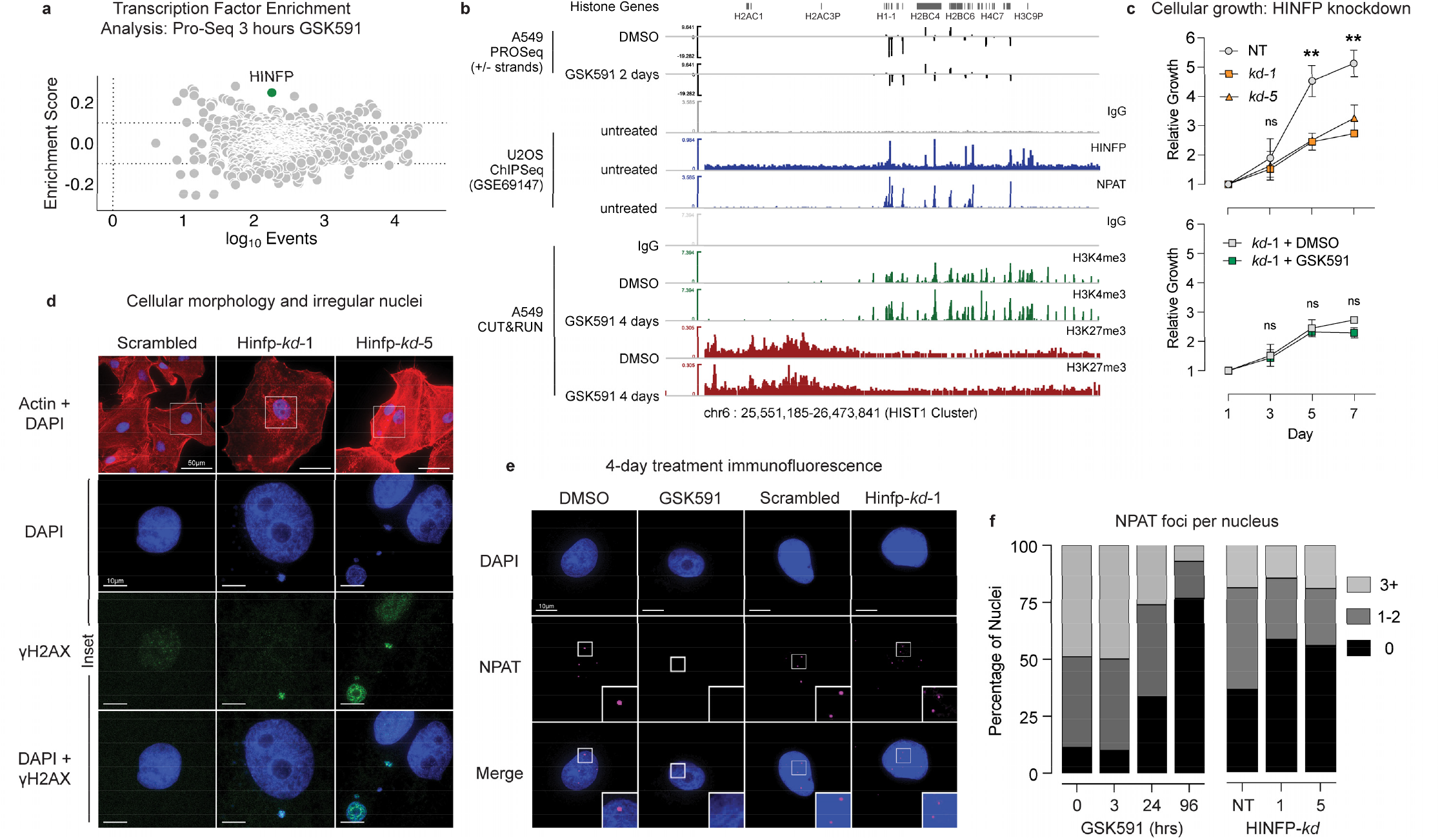
PRMT5 inhibition phenocopies histone transcription arrest caused by HINFP knockdown. **a)** Transcription factor enrichment analysis (TFEA^1^) of PRO-seq data at 3 hours of PRMT5 inhibition by GSK591. **b)** IGV overview plot at the HIST1 locus on chromosome 6: 2-day GSK591 PRO-seq data, U2OS cell ChIP-Seq data for HINFP and NPAT binding (GSE69147^3^), A549 cell H3K4me3 and H3K27me3 CUT&RUN data. **c)** Growth curves measuring metabolic activity of A549 cells after HINFP knockdown ± PRMT5 inhibition; mean ±SD, n=2 biological replicates. Student’s one-tailed t-test; p-value = **<0.005, n.s. = not significant. **d)** A549 cells after 7-days HINFP knockdown, highlighting enlarged “fried-egg” appearance of DAPI-stained nuclei (blue) in actin (red). Immunofluorescence staining of γH2AX (green) overlaid upon nuclei (blue), highlighting γH2AX-positive micronuclei separate from the primary nucleus. Scale bar 50μm in Actin+DAPI and 10μm in inset images. **e)** IF of NPAT in 4-day PRMT5 inhibited and 4-day HINFP knockdown A549 cells. **f)** Count of NPAT foci per nucleus following 0-, 3-, 24-, or 96-hours of PRMT5 inhibition or HINFP knockdown for 4-days; n=1 biological replicate with >200 nuclei counted/condition for GSK591 time course; n=2 biological replicates with >75 nuclei counted/condition for HINFP knockdown.

To assess PRMT5-dependent changes in chromatin state at histone gene loci, we performed CUT&RUN profiling of H3K4me3 and H3K27me3 in cells treated with GSK591 for four days. Despite the observed transcriptional repression of histone genes, the repressive H3K27me3 mark did not spread into the histone gene clusters, and active H3K4me3 marks remained sharply defined (**Figure 3b, bottom**). This absence of heterochromatin spreading suggested that PRMT5 inhibition does not lead to stable chromatin silencing of histone genes. We inferred that transcriptional downregulation likely reflects a dynamic regulatory mechanism rather than chromatin-based gene repression.

To test whether HINFP loss phenocopied PRMT5 inhibition, we performed HINFP knockdown in A549 cells using a lentiviral CRISPRi system^40^. Knockdown efficiency was confirmed by RT-qPCR (**Supplementary Figure 3a**). By immunoblotting total protein lysate after 4-day HINFP knockdown, we observed reduced cellular levels of all core histone proteins and a drastic loss of linker histones (**Supplementary Figure 3b**). Consistent with a cellular stress response to histone depletion, and like PRMT5 inhibition, HINFP knockdown also led to upregulation of p21 (**Supplementary Figure 3a,b**). HINFP knockdown induced growth arrest, beginning around three days after knockdown when the cells had adopted an enlarged “fried egg” appearance (**Figure 3c,d**). Notably, growth arrest following HINFP knockdown was not exacerbated by the addition of PRMT5 inhibition, suggesting that PRMT5 and HINFP may function within a shared pathway to regulate cellular proliferation (**Figure 3c**).

As the enlarged cell morphology closely resembled the phenotype observed with PRMT5 inhibition, we tested these cells for abnormal nuclear structures. Seven days of HINFP knockdown with two separate guide RNA induced ∼45 and 55% of cells to contain multi-irregular nuclei, comparable to the ∼40% observed after 7 days of PRMT5 inhibition (**Figure 3d and Supplementary Figure S3c**). These nuclear abnormalities were consistent with previous reports of aberrant nuclei and genomic instability following HINFP depletion^43^.

Both GSK591 treatment and HINFP knockdown for 4-days reduced the number of immunofluorescence-observed NPAT foci, corresponding with histone locus bodies (HLBs) (**Figure 3e,f**). GSK591 treatment caused a more pronounced effect, depleting both the HLB number and the intensity and size of the HLB that remained (**Supplementary Figure S3d,e**). The loss of HLBs was consistent with 7-day PRMT5 inhibition by both GSK591 and LLY-283 (**Supplementary Figure S3f**).

Given the similarity of these phenotypic consequences, we considered whether HINFP might be a direct substrate of PRMT5. However, HINFP was not found in multiple PRMT5 substrate screens^24,44-46^and its amino acid sequence does not contain any known PRMT5 methylation motifs^24^. E2F1 is reportedly arginine-methylated and is a transcriptional regulator of S-phase progression^47^. However, complete E2F1 knockdown in A549 cells had minimal impact on cellular growth, and PRMT5 inhibition continued to suppress proliferation in the absence of E2F1 (**Supplementary Figure 3g-i**), indicating that E2F1 is not the primary mediator of the PRMT5-inhibition phenotype. Notably, we did not observe E2F1 methylation in our PTMScan studies in A549 cells^24^. We concluded that E2F1 was not the primary mediator of PRMT5-regulated histone gene expression.

Together, these findings demonstrate that both PRMT5 inhibition and HINFP knockdown impair histone gene expression, supporting a shared functional role in regulating histone production. However, their regulatory mechanisms are likely distinct.

### PRMT5 inhibition arrests histone transcription independently of p53 activation

Given our observation of early alternative splicing of MDM4 upon PRMT5 inhibition, we hypothesized that subsequent p53 activation may rapidly induce cellular arrest, thereby altering histone transcription. To test this, we generated *TP53* knockdown cells (**Supplementary Figure S4a,b**), which showed no baseline growth defect on A549 cells (**Figure 4a**). Upon PRMT5 inhibition, *TP53* knockdown cells still exhibited profound growth arrest; further, scrambled control cells exhibited equivalent growth arrest to knockdown (**Figure 4a and Supplementary S4c**), indicating that the antiproliferative effect of PRMT5 inhibition is p53-independent. Moreover, p53-deficient cells developed the same enlarged “fried egg” morphology seen in controls after four days of PRMT5 inhibition, and the multi-irregular nuclear phenotype was not only maintained, but exacerbated, consistent with a role for p53 in suppressing nuclear instability upon PRMT5 loss (**Figures 4b and 4c**). A549 cells treated with a combined p53 knockdown and PRMT5 inhibition were strongly γH2AX-positive by immunofluorescence staining (**Figure 4b and Supplementary Figure S4d**). Although growth arrest persisted in the absence of p53, β-galactosidase staining as a metric for senescence induction was p53-dependent, a result also recapitulated in the non-transformed, *TP53*-wildtype, lung epithelial cell line BEAS-2B (**Figure 4d and Supplementary Figure S4e-f**). These data show that some cellular consequences of PRMT5 inhibition are p53-mediated.

**Figure 4.**
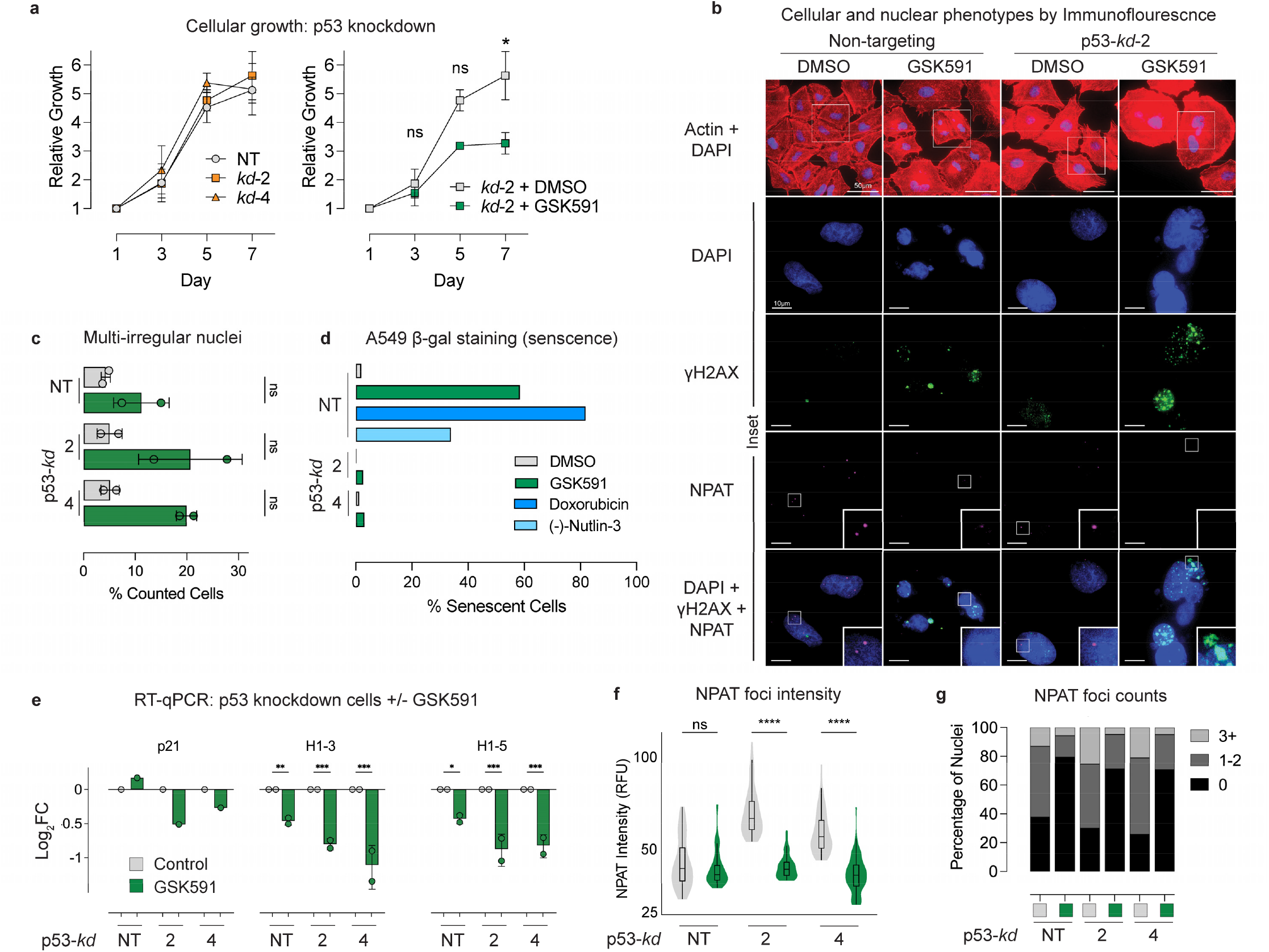
PRMT5 activity is required for histone transcription irrespective of p53 status. **a)** Growth curve measuring metabolic activity of A549 cells after p53 knockdown, ± PRMT5 inhibition; mean ±SD, n=2 biological replicates. **b)** Rhodamine**-**phalloidin (red) and DAPI (blue) stained p53 knockdown A549 cells treated with 4-days PRMT5 inhibition. Immunofluorescence staining of γH2AX (green) overlaid upon DAPI-stained nuclei; NPAT (magenta) labeling histone locus bodies. **c)** Quantified multi-irregular nuclei from p53 knockdown A549 cells ±PRMT5 inhibition; n=2 biological replicates, >125 cells counted/replicate. **d)** β-galactosidase staining positivity of p53 knockdown cells ±PRMT5 inhibition in A549 cells. **e)** RT-qPCR analysis of p53 knockdown A549 cells treated with 24-hours GSK591. **f-g)** NPAT foci intensity and count ±p53 knockdown ±PRMT5 inhibition quantified from immunofluorescence slides; n=2 biological replicates, >125 cells counted/replicate. Student’s one-tailed t-tests were applied; p-value = *<0.05, **<0.005, ***<0.0005, ****<0.00005, n.s. = not significant.

Finally, we assessed whether histone transcription loss upon PRMT5 inhibition was p53-independent. While p21 induction depended on p53 presence, histone gene expression was reduced after 24 hours of PRMT5 inhibition in p53 knockdown cells, mirroring the response in scrambled controls (**Figure 4e and Supplementary S4g**). Consistent with these findings, IF confirmed that GSK591 treatment reduced HLBs per cell, NPAT intensity at HLBs, and HLB area, regardless of p53 status (**Figure 4b,f,g and Supplementary Figure S4h**).

Together, these results indicate that PRMT5 promotes cellular proliferation, maintaining histone transcription and nuclear integrity through a p53-independent mechanism.

### PRMT5 activity maintains histone transcription throughout S phase

We hypothesized that PRMT5 activity is required to initiate histone transcription at the onset of S phase. To test this, we arrested cells in G1 phase of the cell cycle with the CDK4/6 inhibitor palbociclib^48^and released into S-phase by exchange of media containing palbociclib with fresh media, with or without PRMT5 inhibition (**Figure 5a**). Over the course of 11 hours post-release from palbociclib arrest, transcript markers of G1/S phase transition (Cyclin E1) decreased, the S phase marker PCNA rapidly increased and peaked at 7-hours post release, and the G2 marker Cyclin A2 increased across the time series (**Supplementary Figure S5a**). PRMT5 inhibition did not impair early S-phase entry; S-phase transcripts including Cyclin E, PCNA, and E2F1 increased rapidly and peaked between 3-and 7-hours post-release in both control and PRMT5-inhibited cells; Cyclin A2 expression rose steadily throughout S phase, consistent with progression toward G2/M (**Supplementary Figure S5b**). While histone RNA levels were indistinguishable between control and PRMT5-inhibited cells during early S phase, by 9 hours post-palbociclib release, histone transcript levels were reduced in PRMT5-inhibited compared with control cells (**Figure 5b and Supplementary Figure S5b**). These data show that PRMT5 activity is dispensable for initiating S phase and histone transcription but is required to sustain transcriptional activity throughout S phase.

**Figure 5.**
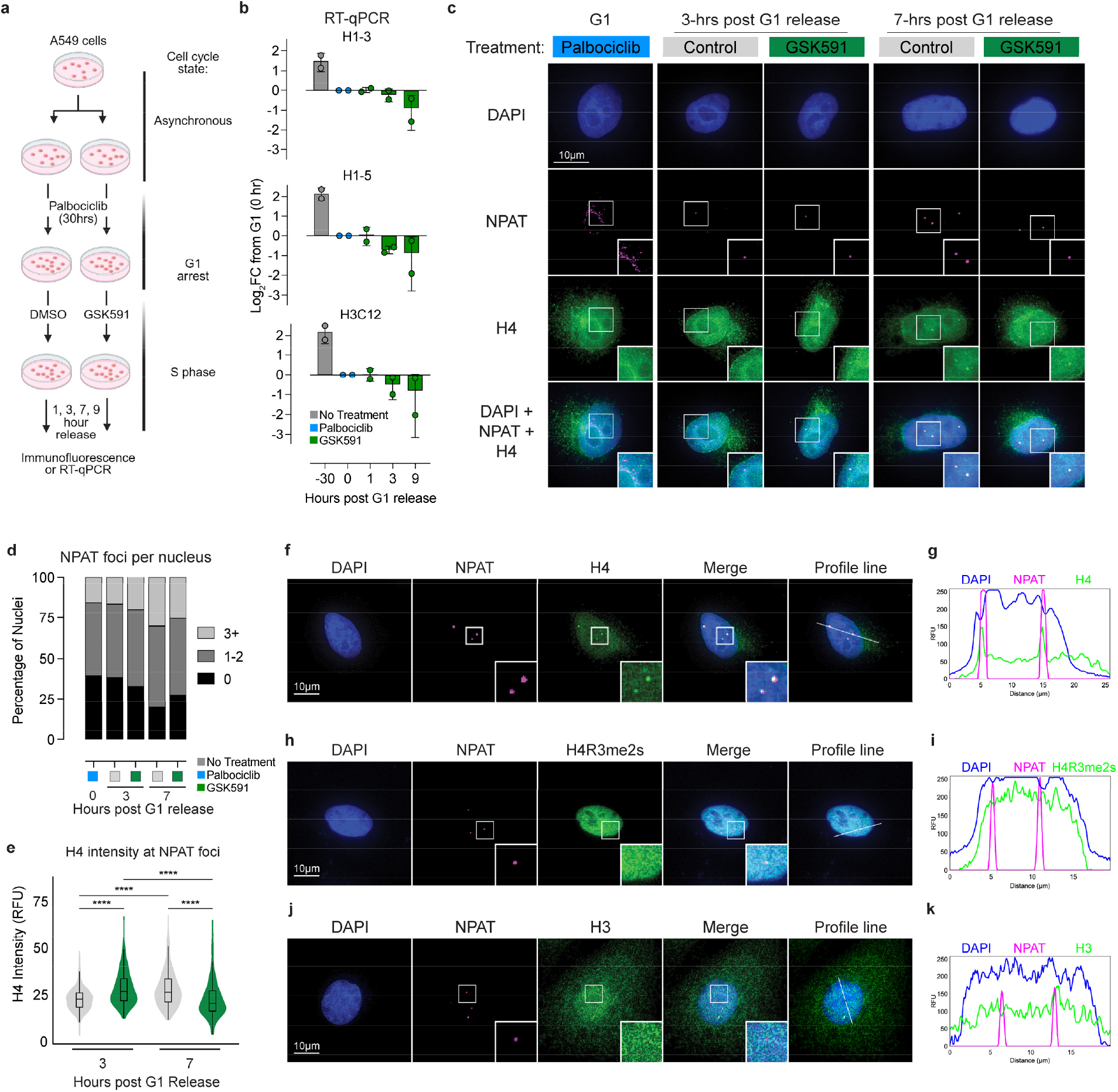
PRMT5 activity maintains histone transcription in S phase. **a)** Schematic of palbociclib G1 arrest with release into S phase. **b)** RT-qPCR of representative histone transcripts throughout S phase ± PRMT5 inhibition. **c)** IF throughout S phase ± PRMT5 inhibition of NPAT visualizing NPAT foci (histone locus bodies; magenta) and histone H4 (green) overlaid upon DAPI-stained nuclei (blue). **d-e)** NPAT foci count and intensity throughout S phase ± PRMT5 inhibition; n=1 biological replicate, >100 cells/condition. Student’s one-tailed t-tests was applied; p-value = ****<0.00005, n.s. = not significant. **f-g)** IF of NPAT (magenta), H4 (green), and DAPI-stained nuclei (blue) and profile line plot of intensities, highlighting overlap of NPAT and H4. **h-i)** IF of NPAT (magenta), H4Rme2s (green), and DAPI-stained nuclei (blue) and profile line plot of intensities, highlighting lack of overlap between NPAT and H4Rme2s. **j-k)** IF of NPAT (magenta), H3 (green), and DAPI-stained nuclei (blue) and profile line plot of intensities, highlighting lack of overlap between NPAT and H3.

At the onset of S phase, NPAT phosphorylation by CDK2/Cyclin E initiates the formation of NPAT foci (defined as HLBs) to recruit transcriptional and pre-mRNA processing machinery and facilitate histone transcription in S phase. Consistent with altered histone transcription in S phase, PRMT5-inhibited cells failed to increase the number of NPAT foci throughout S phase, indicating either failure to properly form histone locus bodies or disruption of their integrity (**Figure 5c,d**). While the NPAT puncta maximal-projection area was similar between conditions, NPAT intensity was greater at 3-hours into S phase with PRMT5 inhibition (**Supplementary Figure S5c,d**). We therefore concluded that PRMT5 activity is necessary in S-phase to maintain normal HLBs.

A recent preprint reported that soluble histone H4 protein is present at histone locus bodies and may exert negative feedback on histone transcription in both *D. melanogaster* and human cells^49^. We and others previously showed that histone H4 Arginine 3 (H4R3) is a robust substrate of PRMT5 *in vitro* (H4R3me2s), but the biological role(s) of H4R3me2s in cells remains unclear^50, 51^. Importantly, H4 is not a PRMT5 substrate when assembled into nucleosomes^50, 52^. Therefore, we hypothesized that the role of PRMT5 activity in regulating histone transcription in S phase might be mediated by methylation of soluble H4 protein. To test this hypothesis, we repeated the palbociclib release experiments and then probed for H4 protein accumulation at NPAT foci. Validation immunoblots for the H4 antibody confirmed its selectivity (**Supplementary Figure 5e,f**). We observed a greater increase of H4 intensity at NPAT puncta at 3-hours post-release into S phase in PRMT5-inhibited cells compared to control cells (**Figure 5e**). Consistent with the Ahmad *et al*.^49^model of H4 negative feedback on histone transcription at the end of S phase within HLBs, we observed an accumulation of H4 signal at control HLBs over the period of 3 to 7-hours postrelease, while PRMT5-inhibited cells peaked H4 intensity early and diminished H4 intensity by 7-hours (**Figure 5e**).

Histone H4 is generally associated with histone H3 in dimers and tetramers or in a nucleosome with all four core histones^5^. To test if histone H4 foci are uniquely found with NPAT puncta, we tested asynchronous cells by IF for both H4 and H3 protein localization. H4 immunofluorescence revealed pan-nuclear staining—and some perinuclear staining—with local foci which overlapped with NPAT signal (**Figures f,g**). We hypothesized that PRMT5 methylation of H4R3 might influence this localization; strikingly, we did not observe any NPAT-associated nuclear foci with an H4R3me2s antibody (**Figure 5h,i**). These data are consistent with the hypothesis that PRMT5 methylation of H4R3 antagonizes premature H4 negative feedback by limiting H4 accumulation at the HLB to sustain histone transcription throughout S phase. Notably, H3 staining did not exhibit any enrichment at NPAT foci (**Figures 5j,k**), supporting the notion that H4 monomers are accumulated at the HLB. Further, we observed detergent-extracted soluble H4—but not other core histones—was lost over 4- and 7-days of PRMT5 inhibition (**Supplementary Figure 5g**). Finally, we tested if PRMT5 accumulated at NPAT foci, but we did not observe any selective enrichment (**Supplementary Figure 5h,i**).

These data are consistent with a hypothesis that PRMT5 methylation of H4R3 antagonizes premature H4 negative feedback by limiting H4 accumulation at the HLB to sustain histone transcription throughout the S phase.

### PRMT5 activity is required for histone protein production during S phase

As we established that PRMT5 activity reduces histone gene transcription within the first S-phase following inhibition, we questioned whether loss of transcription resulted in altered histone protein levels within the cell. We acid-extracted chromatin-bound proteins from asynchronous A549 cells treated with 4-days of PRMT5 inhibition and observed markedly reduced levels of both core and linker histones on immunoblot, as normalized by live cell count prior to extraction (**Figure 6a**). This reduction in histone protein was consistent across multiple experimental replicates and was observed with both 7-day PRMT5 and HINFP knockdowns (**Figure 6a and Supplementary Figure S6a**). Additionally, knockdown of PRMT5 in both K562 cells using CRISPRi and mouse embryonic stem cells (mESCs) using shRNA also exhibited cell-count normalized loss of histone proteins (**Figure 6a and Supplementary Figure S6b**). As sample input was normalized by cell number, this consequence was not attributable to changes in total cell number or viability. As we previously reported^53^, both PRMT5 inhibition and knockdown resulted in increased chromatin-bound SNRPB spliceosomal protein. In contrast, HINFP knockdown did not exhibit increased chromatin-bound SNRPB (**Supplementary Figure S6a**). To further characterize histone protein loss upon PRMT5 knockdown, we completed HPLC analysis of acid-extracted histones. For both K562 cells and mESCs, H1 linker histones were depleted with respect to core histones (**Supplementary Figure S6c**).

**Figure 6.**
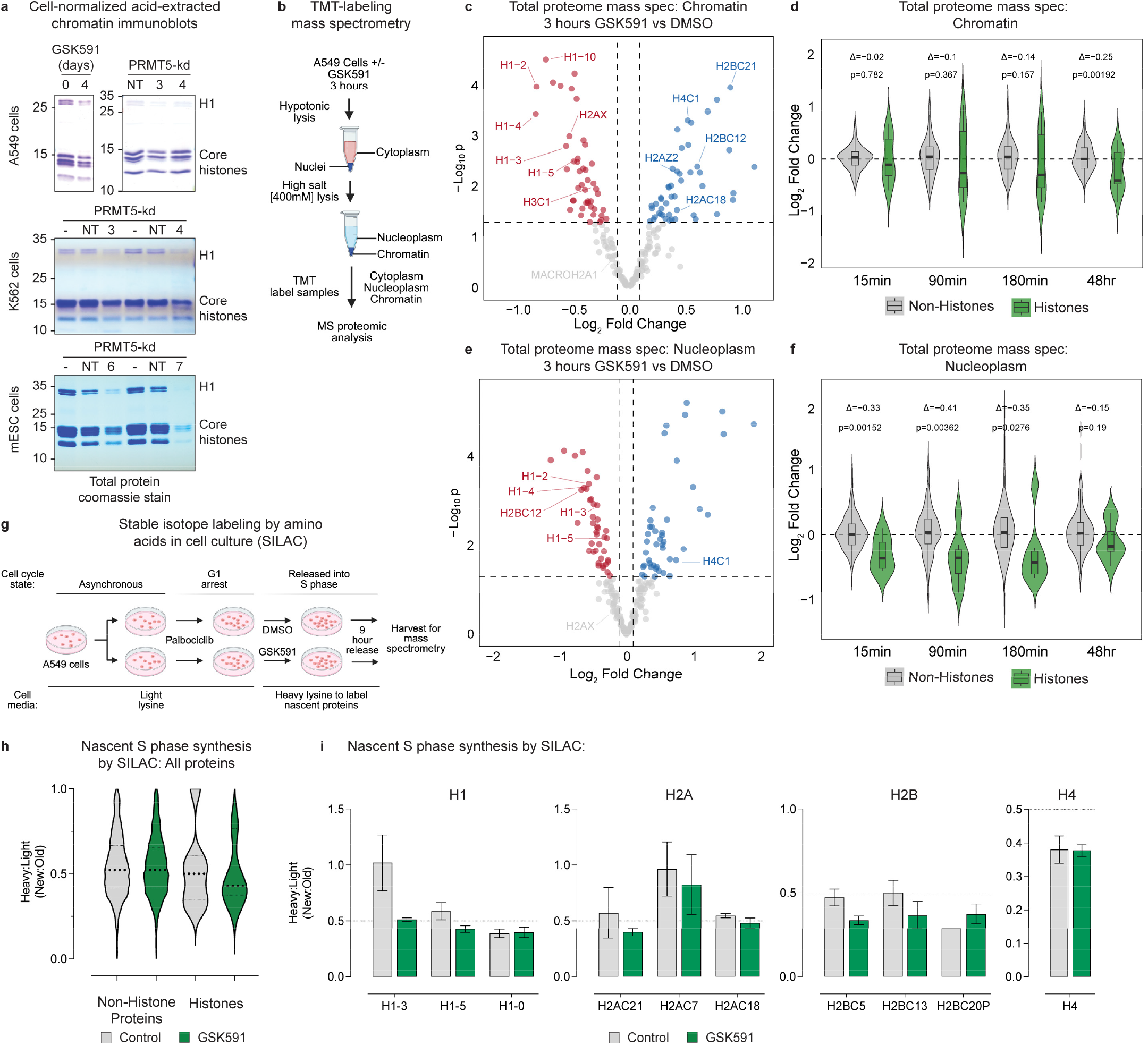
PRMT5 activity is required for histone protein production during S phase. **a)** Immunoblots of acid-extracted chromatin from equal cell number; top: 4 days of PRMT5-inhibition or 7 days of PRMT5 knockdown in A549 cells, middle: 5 days of PRMT5 knockdown in K562 cells, bottom: 4 days of PRMT5 knockdown in mESCs (un-transfected negative control (-) and non-targeting negative control (NT)). **b)** TMT-labeling mass spectrometry experimental set-up. **c)** Total proteome mass spectrometry on the chromatin fraction. **d)** Non-histone vs histone Log_2_FC over time; chromatin fraction. Random permutation test (10,000). **e)** Total proteome mass spectrometry on the nucleoplasmic fraction. **f)** Non-histone vs histone Log_2_FC over time; nucleoplasmic fraction. Random permutation test (10,000). **g)** Experimental set-up for G1 phase-release SILAC mass spectrometry. **h)** Nascent S phase synthesis by SILAC; histones vs non-histones. **i)** Nascent S phase synthesis for selected histones. Bars represent mean ±SEM of 4 replicates.

To quantitatively test if histone proteins were reduced from chromatin upon PRMT5 inhibition, we treated A549 cells for 15min, 1.5-, 3-, and 48-hours and completed fractionated, tandem mass tag (TMT) quantitative proteomics on cytoplasm, nucleoplasm, and chromatin (**Figure 6b, Supplementary Figure S6d,e, and Supplementary Table S3**). After 3 hours of PRMT5 inhibition, we observed a loss of multiple linker histones and H3 from chromatin, with an increase of H2B and H4 (**Figure 6c**). Comparing histone abundance on chromatin between PRMT5-inhibited vs control over the entire experiment, we observed progressive loss of histones from chromatin over time points of PRMT5 inhibition (**Figure 6d and Supplementary Figure S6f**). These findings were reflected in the nucleoplasm as well, with linker histones reduced in abundance in nucleoplasm at 3 hours and beyond (**Figure 6e,f**). Notably, H4 was the sole histone increased in abundance both the nucleoplasm and cytoplasm following PRMT5 inhibition (**Figure 6e and Supplementary Figure S6g**). By 48-hours, there was a significant increase in histone proteins in the cytoplasm of PRMT5-inhibited cells vs control cells (**Supplementary Figure S6h**). These experiments quantitatively demonstrated that PRMT5 inhibition reduced total histone levels in nucleoplasm and on chromatin within 3 hours of treatment, suggesting that histone deposition is dependent upon PRMT5 activity.

We next asked if nascent histone production was being altered by PRMT5 inhibition. To directly test if histone synthesis during S phase relied on PRMT5 activity, we employed stabile isotopy labeling with amino acids in cell culture (SILAC)-based metabolic labeling: cells were first grown and arrested at G1 phase in lysine-depleted media with added light lysine, then released into S phase in lysine-depleted media with heavy [^13^C_6_, ^15^N_2_]-lysine to label nascent proteins (**Figure 6g, Supplementary Table S4**). Quantitative (heavy/light ratio) proteomic analysis of samples from 9 hours post release revealed a reduction in some newly synthesized histone proteins in PRMT5-inhibited cells compared to controls, whereas non-histone proteins were not impacted (**Figure 6h**). We observed that PRMT5 inhibition specifically reduced the nascent production of some linker histone H1 isoforms while increasing that of others (**Figure 6i and Supplementary Figure S6i**). Both H2A and H2B histones had a downward trend in nascent synthesis in PRMT5-inhibited cells (**Figure 6i**). Nascent histone H3 protein production appeared increased with PRMT5 inhibition, while H4 and the production of other histone variants were unchanged in the two conditions (**Supplementary Figure S6i**). Together, these data support the conclusion that PRMT5 activity both regulates histone transcription and histone protein production within S phase.

### Histone H4 negatively regulates feedback on histone transcription to induce micronuclei

Given the requirement for PRMT5 to sustain histone transcription during S phase, we hypothesized that it may act directly through methylation of a critical histone transcription regulator. If soluble histone H4 protein accumulates at NPAT puncta to mediate a role as a sensor to downregulate histone transcription at the end of S phase^19^, we hypothesized that PRMT5-dependent methylation of H4 could influence this feedback mechanism. To directly assess the role of H4R3 methylation in this process—with the hypothesis that negative feedback at the HLB requires wild type (wt) H4R3—we overexpressed C-terminally HA-tagged H4-wt, the methylation-deficient mutants H4R3K and H4R3A, and the oncohistone mutant H4R3C^54^(**Figure 7a**). Overexpression at 4-days was confirmed by HA immunoblot, with comparable protein levels between mutants observed (**Supplementary Figure S7a**). Probing the HA tag by IF, we observed that all overexpressed histones localized exclusively to the nucleus and exhibited enrichment in nuclear foci overlapping with NPAT (**Figure 7b**). There was minimal HA signal in GFP control cells (**Supplementary Figure S7b**).

**Figure 7.**
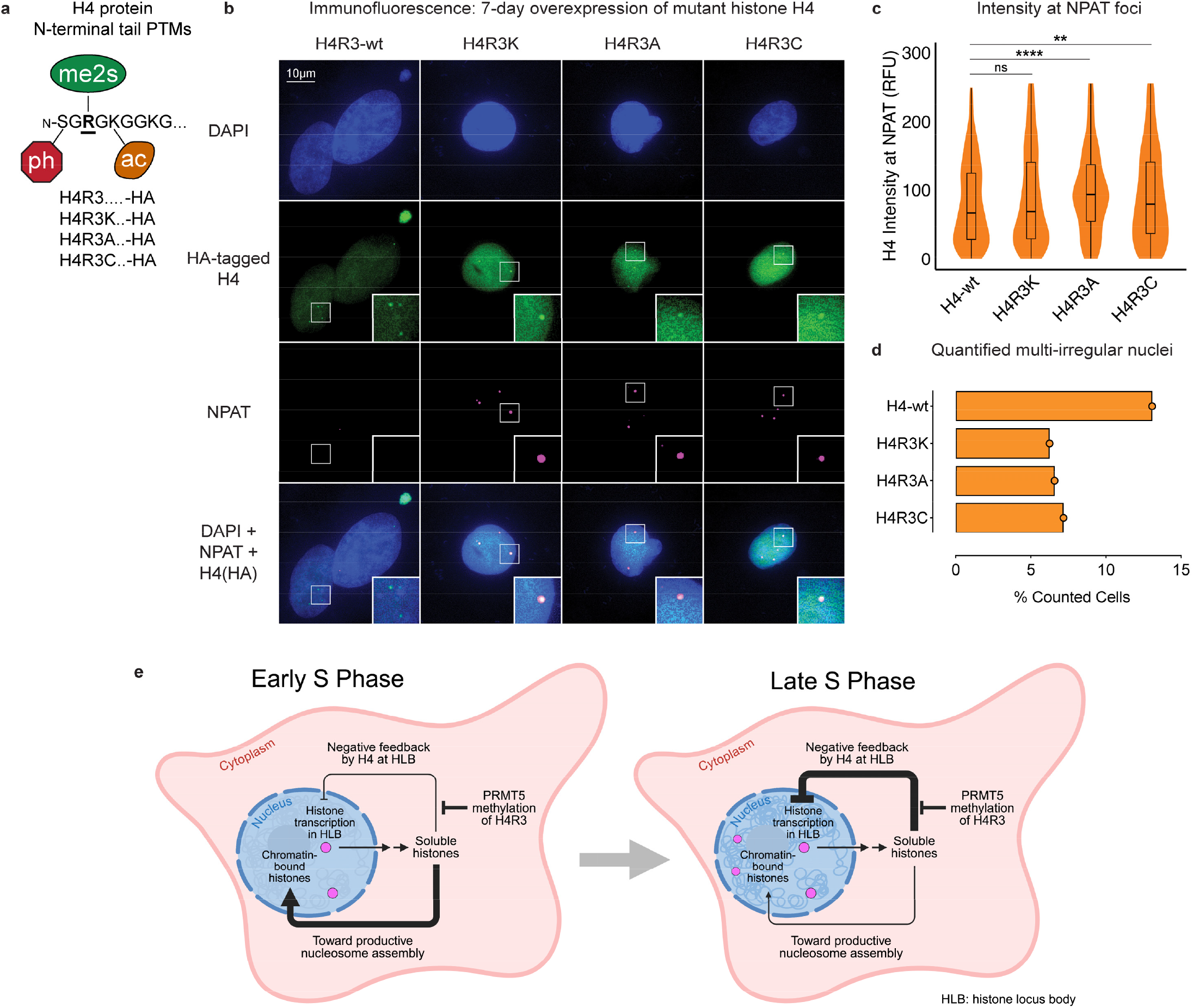
Histone H4 negatively regulates feedback on histone transcription to induce micronuclei. **a)** Representative H4 N-terminal tail highlighting H4S1phosphorylation, H4R3 symmetric dimethylation, and H4K5acetylation. **b)** Immunofluorescence of overexpressed HA-tagged H4 in A549 cells. **c)** Intensity of HA-H4 at NPAT foci; n=1 biological replicate, >225 cells/condition. **d)** Count of multi-irregular nuclei per cell; n=1 biological replicate, >225 cells/condition. **e)** A unifying hypothesis on histone transcription regulation by PRMT5: PRMT5 methylates nascent histone H4 in S phase to prevent premature H4 accumulation at histone locus bodies, delaying transcriptional feedback to support histone production. Created in BioRender.

Consistent with our findings with PRMT5 inhibition, overexpression of H4-wt increased micronuclei formation in cells whereas the methylation-deficient mutants did not increase micronuclei, even though the H4R3A and H4R3C presence at NPAT-positive foci was increased compared to H4-wt (**Figure 7c,d**). Overexpression of H4-wt and mutant H4 had little impact on NPAT foci area or count, with mutant H4 significantly increasing NPAT foci intensity compared to H4-wt (**Supplementary Figure S7c-e**). Overall, these observations support the hypothesis that H4 dynamics at HLBs—as mediated by PRMT5 activity towards H4R3—participate in regulating the duration of histone gene expression and the production of histone proteins during S phase (**Figure 7e**).

## Discussion

As a unifying explanation for PRMT5’s critical cellular function has remained elusive, here we set out to understand the immediate cellular consequences of PRMT5 inhibition. We demonstrated that histone gene transcription is a direct and essential function of PRMT5 activity and is required to prevent subsequent loss of histone proteins and genome integrity. Furthermore, our data supports the model in which symmetric dimethylation of histone H4 at arginine 3 (H4R3me2s) controls negative feedback by H4 protein on histone transcription. As there is widespread interest in approaches to target PRMT5 for cancer chemotherapy with multiple clinical trials underway, understanding PRMT5’s essential function is necessary for patient stratification and combination therapies. Overall, our findings establish PRMT5 as a central regulator of histone homeostasis and genome stability, providing a mechanistic explanation for its essential role in proliferating cells.

Our data show that previously described PRMT5 functions, such as regulation of splicing and p53 signaling ^34-36, 55-57^, are insufficient to account for its critical role in sustaining proliferation. While a prior study suggested that p53 status is important for PRMT5 inhibitor sensitivity^58^, our findings show that histone transcription defects and proliferation arrest persist in p53-deficient cells, demonstrating a p53-independent mechanism. Remarkably, nascent transcription of histone genes is lost within three hours of PRMT5 inhibition, preceding detectable loss of symmetric dimethylarginine. This rapid transcriptional repression suggests that PRMT5 acts on a labile, S phase–specific substrate.

Prior studies characterized multiple PRMT5 substrates involved in cell cycle regulation, including E2F1^47^and others^59^, but few addressed PRMT5’s role at specific cell cycle stages or considered its rapid transcriptional effects. Given that histone H4 Arginine 3 is a canonical PRMT5 substrate^60-64^and that recent work suggests H4 may exert feedback on histone transcription^49^, we hypothesized that H4 could be the critical mediator of this function. Furthermore, as we and others previously showed that PRMT5 cannot methylate histones within nucleosomes^50, 52^, we hypothesized that methylation occurs on soluble H4 in cells. In this model, unmethylated H4 accumulates at the histone locus body (HLB) to mediate transcriptional feedback and termination at the end of S phase. PRMT5-mediated methylation may delay this feedback by limiting H4’s accumulation or interaction with transcriptional machinery. Supporting this model, our data show that upon PRMT5 inhibition, H4 accumulates prematurely at HLBs and that methylation-deficient H4 mutants localize more robustly to HLBs than wildtype H4.

Other approaches to histone depletion in mammalian systems, including knockdown of HINFP^43^or CASP8AP2 (also known as FLASH)^3^, produce similar outcomes to loss of PRMT5 activity: reduced histone transcription, slowed proliferation, and chromatin abnormalities without immediate cell death. In each case, cells continue to synthesize DNA but accumulate replication stress. Indeed, linker histones are lost to a greater extent than core histones after PRMT5 knock-down, consistent with H1 as genome protective protein. We observed a reduction in total cellular DNA levels after PRMT5 inhibition; however, cells did not die. Consistent with a premature arrest of histone production in late S phase, we hypothesize that non-essential late-replicating DNA is lost. Future studies will be necessary to test this hypothesis. While PRMT5 is frequently overexpressed in tumors, here we show that its essentiality stems from maintaining histone production, rather than more general transcriptional or splicing roles. This highlights a new therapeutic vulnerability of uncoupling histone synthesis from DNA replication. While additional studies are necessary, we speculate that PRMT5 overexpression may enhance histone output to support rapid tumor cell division, providing a rationale for its upregulation in many cancers.

Consistent with its potential restriction to specific cell cycle windows, quantification of H4 Arginine 3 (H4R3me2s) in cells suggests it is a rare mark^65-67^, and no reliable study exists demonstrating its enrichment in transcriptional regulatory epigenomic loci. As we showed for soluble H2A Arginine 3 symmetric dimethylation in *Xenopus* eggs and embryos^68^, we propose that H4R3 symmetric dimethylation serves as a pre-deposition modification that couples S phase progression with histone supply, rather than as canonical transcription-regulatory chromatin modification. Notably, the low micromolar Km (Michaelis constant) of PRMT5 for its methyl donor S-adenosylmethionine (SAM) is comparable to intracellular SAM concentrations, suggesting PRMT5 activity could be sensitive to metabolic flux. This raises the possibility that PRMT5 integrates methionine and one-carbon metabolism with cell cycle progression to modulate histone production. Overall, our model aligns with emerging views that histone proteins themselves can act as sensors in feedback circuits regulating their own expression^20, 69^.

Histone H4 is one of the most evolutionarily conserved proteins across eukaryotes, with nearly identical sequences from yeast to humans^70, 71^. In contrast to other core histones, which are encoded by multiple variant genes clustered in the HIST1 and HIST2 loci, all canonical H4 genes encode the same protein sequence and lack known variants^16^. This high conservation underscores a potentially unique regulatory role for H4 in chromatin biology. Notably, somatic “oncohistone” mutations in H4 have been identified in cancer, including a recurrent R3C mutation^54^. In our study, overexpression of H4R3C resulted in fewer micronuclei than wild-type H4, suggesting this mutation may blunt the negative feedback response and promote continued DNA synthesis. Further investigation will be needed to determine whether such mutations help cancer cells tolerate chromatin stress by decoupling histone supply and transcriptional control.

In summary, we propose that PRMT5 promotes proliferation by sustaining histone gene transcription through a feedback circuit involving H4 methylation. Loss of PRMT5 disrupts this balance, leading to reduced histone supply, chromatin instability, and genome fragmentation. This model provides a mechanistic explanation for PRMT5’s essentiality and offers new insight into how cells coordinate chromatin production with DNA replication.

### Limitations and future directions

While our findings establish PRMT5 as a critical regulator of histone transcription and chromatin integrity during S phase, several questions remain. The precise mechanism by which PRMT5 sustains histone transcription—whether through direct chromatin engagement, methylation of histone H4R3, methylation of transcriptional cofactors, or altered histone mRNA or protein stability—requires further investigation. In addition, although we observe premature termination of histone gene expression upon PRMT5 inhibition, coincident with H4 accumulation at the HLB, the cellular mechanism linking this observation to histone transcription arrest remains undefined. PRMT5 has many protein substrates and regulates multiple biological processes; future studies are required to determine whether histone insufficiency alone is sufficient to induce genome destabilization or whether other PRMT5-regulated processes contribute to the observed phenotypes. Future studies will explore additional PRMT5 substrates as candidate mediators—for example LSm11^72^, probe the interactome of methylated and unmethylated H4 using mutant proteins, and employ temporal degron overexpression systems to precisely express these histones in S phase to resolve these mechanisms.

## Methods

### Cell Culture

A549 (CCL-185), BEAS-2B (CRL-9609), and K562 (CCL-243) cells were purchased from ATCC. A549 and BEAS-2B were cultured in Dulbecco’s Modified Eagle Medium (DMEM; Corning: 10-013-CV) supplemented with 10% FBS (Hyclone), 100 μg/ml streptomycin and 100 IU/ml penicillin (Corning). K562 cells were cultured in Iscove’s Modified Dulbecco’s Medium (IMDM; Gibco) supplemented with 10% Benchmark fetal bovine serum (FBS) and 1x penicillin-streptomycin. Mouse embryonic stem cells (mESCs) were prepared in house^73^and cultured in high-glucose Dulbecco’s Modified Eagle Medium (DMEM; Gibco: 11965092), supplemented with 15% STASIS fetal bovine serum (FBS), 1x non-essential amino acids (NEAA), 1 mM sodium pyruvate, 2 mM L-glutamine, 1x penicillin-streptomycin, 0.1 mM β-mercaptoethanol, and leukemia inhibitory factor (LIF; prepared in-house by the core facility).

All cells were maintained at 37°C in 5% CO_2_ with humidity. Cell identity was confirmed by STR profiling and *Mycoplasma* negativity was confirmed by PCR (primers in **Supplemental Table S5**)^74^. All compounds except for blasticidin (water) were dissolved in DMSO and were added to cell media at a maximum of 0.01% final volume.

### Immunoblots

Whole cell lysis was completed by 30-minute incubation in RIPA buffer (1% NP-40, 150 mM NaCl, 1mM EDTA, 50 mM Tris-HCl pH 8 at 4°C, 0.25% Sodium Deoxycholate, 0.1% SDS, 1x protease inhibitor, and 1x phosphatase inhibitor) at 4°C with rotation. Samples were probetip sonicated 3×5sec at 20% amplitude to liberate chromatin-bound proteins and lysate was cleared of insoluble material. Immunoblots were performed on PVDF (Immobilon, Millipore) and detected by secondary antibodies at 1:1000 (Sigma: NA934 or NA931) and ECL chemiluminescence (Lumigen: TMA-6), imaged with an ImageQuant LAS4000 (GE). Primary antibodies are reported in **Supplemental Table S5**.

### Acid extraction of chromatin to isolate histones

This protocol was adapted from our previously published protocol^75^. Briefly, cell pellets (cell count normalized where indicated) were re-suspended in hypotonic lysis buffer (HLB: 10 mM Tris-HCl pH 8.0, 1 mM KCl, 1.5 mM MgCl2, and 1 mM DTT, with protease and phosphatase inhibitors) and incubated for 30 minutes on rotation at 4°C to promote hypotonic swelling and mechanical lysis, followed by centrifugation to pellet intact nuclei. Nuclei were rinsed once in HLB then resuspended in nuclear lysis buffer (NLB: 10 mM Tris-HCl pH 8.0, 250 mM KCl, 0.1% NP40, and 1 mM DTT, with protease and phosphatase inhibitors) and rotated for 30 min at 4°C. Chromatin was then collected via centrifugation and rinsed with NLB. Chromatin was resuspended in 0.4N H_2_SO_4_, rotated for 30 min. at 4°C, then pelleted at 16,000xg for 10 min at 4°C to remove debris. TCA addition to the supernatant precipitated histones, followed by centrifugation and ice-cold acetone washes to collect and rinse the histones. After air-drying, the histone pellet was dissolved in ddH_2_O and either analyzed as is (if pre-normalized by cell number) or scaled by cell-number-matched DNA content as determined by TRIzol extraction. Samples were analyzed by SDS-PAGE gel and Coomassie staining or immunoblotting.

### Indirect immunofluorescence

Cells were seeded on poly-L-lysine coated coverslips (Neuvitro: GG-18-PLL) and grown for a minimum of 4 days in the presence of 0.01% DMSO, 0.5μM or 1μM compound (See **Supplemental Table S5** for dosing) for 1-7 days. Cells were rinsed with warm PBS (Corning) and fixed in 4% methanol-free formaldehyde (ThermoScientific) for 10 minutes at room temperature, rinsed with PBS, and either stored at 4°C in PBS until use or immediately quenched with 0.1M ultraPure glycine (Invitrogen) and blocked 1 hour in 1% fish skin gelatin in PBS with 0.01% Tween-20. Primary antibodies (**Supplemental Table S5**) were prepared in blocking solution and inverted coverslips were incubated on antibodies overnight at 4°C in a humid chamber. Coverslips were rinsed in PBS and incubated with secondary antibody (Thermo: 35552, Invitrogen: A21245 or Invitrogen: A21235) at 1:1000 in blocking buffer containing actin-rhodamine at 1:33 (Invitrogen: R37112) for 1 hour at room temp in a humid chamber protected from light. Coverslips were rinsed in PBS and mounted to glass slides with DAPI prolong gold anti-fade (Thermo: P36930).

Images were acquired with a Plan Apochromatic 60x objective (model BZ-PA60, NA 1.4/0.13 mm) and bandpass filters (Chroma Technology: DAPI: OP-87762/49028-UF1, EGFP: OP-87763/49002-UF1, CY3/TRITC: OP-87764/49004-UF1, CY5: OP-87766/49006-UF1) on a Keyence epi-fluorescence microscope (BZ-X810).

Images for figure visualization were captured as 11-image z-stacks with 0.2μm pitch and presented as a maximal projection. Pseudo-colored images were processed in FIJI (ImageJ2, Version: 2.14.0/1.54f, Build: c89e8500e4) with custom macros to adjust levels equally across all treatments of each experiment^76^. Profile line plots were completed in FIJI. All quantifications were completed on single z-stack images with custom macros in FIJI; macros are shared on https://github.com/Shechterlab/Roth_etal_2025.

### Nuclear morphology analysis

Cells were fixed and prepared for imaging as described above. A cell was scored as mono-nuclear if it contained one nucleus, bi-nucleate if it contained two nuclei of equal size, and multi-irregular if it contained one or more micronuclei, contained >1 nucleus which were of unequal size, or exhibited nuclear bulging. Most cells scored as multi-irregular contained micronuclei. At least 100 cells were scored per condition, totaling >300 cells over three biological replicates for most experiments.

### RNA isolation and RT-qPCR

RNA was purified with the Monarch Total RNA kit (New England Biolabs) and reverse transcribed with Moloney Murine Leukemia Virus (MMLV) reverse transcriptase (Invitrogen) and Random Hexamer primers (Integrated DNA Technologies). Abclonal master mix (RM21203) was used to quantify cDNA on a QuantStudio Pro 6. An initial 95°C for 5 min was followed by 45 cycles of amplification using the following settings: 95°C for 15 seconds, 60°C for 1 minute. The efficiency-corrected threshold cycle (ΔCT) method was used to calculate the relative levels of RNA with expression normalized to vinculin^77^. Melting-curve analysis was performed to ensure primer specificity; primer sequences are reported in **Supplementary Table S5**.

### MTS cellular proliferation assay

Cells were plated in 96-well plates and relative growth rate quantification was completed per the manufacturer’s instructions (Promega: G3580).

### Precision Run-On (PRO) nascent transcriptome sequencing

PRO-seq was performed as described previously^78^. Briefly, approximately 10 million A549 cells were treated in T182 flasks. At the conclusion of the treatment, media was aspirated, and cells were rinsed with 10mL of icecold PBS, scraped, and collected into 15mL conical tubes on ice. Cells were collected by centrifugation at 1,000g at 4°C for 5 minutes. PBS was aspirated and pellets were resuspended in 10mL ice-cold permeabilization buffer. Cells were incubated in permeabilization buffer for five minutes before being centrifuged at 1,000g at 4°C for five minutes. Permeabilization buffer was aspirated and permeabilized cells were resuspended in 100uL freezing buffer and snap frozen before being stored at −80°C. When preparing for the run on step, frozen permeabilized cells were thawed on ice. Run-on master mix was warmed to 37°C and 100uL of 2x run-on master mix was added to 100uL of frozen cells before being placed in a 30°C water bath for five minutes with slight agitation after 2.5 minutes of incubation. After five minutes of run-on, 500uL of Trizol LS was added and samples were vortexed until homogenized to stop the reaction. 130uL of chloroform was added, samples were vortexed for 15 seconds, and tubes were centrifuged at 15,000g for five minutes at 4°C. The aqueous phase was transferred to a new Eppendorf tube and RNA was precipitated using 2.5 volumes of 100% ethanol for 10 minutes before being centrifuged at 15,000g for 20 minutes at 4°C. The pellet was washed with 75% EtoH, centrifuged again, and ethanol removed completely before resuspending the RNA pellet in 19uL of DEPC-treated water. Sample quality was assessed by nanodrop and Tapestation before proceeding to RNA fragmentation with Ambion RNA fragmentation reagent (Ambion: AM8740) for 4 minutes at 72°C before adding stop solution to the reaction. Free NTPs were removed using a p30 column. Nascent RNA was enriched using streptavidin-coated Dynabeads (Thermo Fisher: 65305), washed 2X with 500uL of ice cold high-salt wash buffer using a magnet. Beads were then washed 2X with 500uL binding buffer and 1X with low-salt wash buffer. Buffer was removed and beads were resuspended in 300uL of Trizol to dissociate nascent RNA from the beads. All buffers were made according to Mahat, 2016. 60uL of chloroform was added and vortexed. The mixture was centrifuged at 15,000g for five minutes at 4°C. The aqueous phase was removed and moved to a separate tube before repeating the nascent RNA enrichment and wash steps. After the second round of purification, nascent RNA was precipitated with 1.25uL glycoblue and 2.5 volumes of 100% ethanol before centrifugation at 15,000g for 20 minutes at 4°C. Ethanol was removed and the RNA pellet was washed with 75% ethanol. RNA was centrifuged again before removing the 75% ethanol completely. Pellets were resuspended in 5uL DEPC treated H2O and taken directly into library preparation using the NEBNext Ultradirectional II RNA sequencing kit (NEB: E7760S) following the manufacturer’s instructions. Samples were sequenced using an Illumina HiSeq.

PRO-seq data were processed using the PEPPRO pipeline (v1.0)^79^. Genome references including hg38 FASTA, Bowtie2 indices, GTF annotations, and rDNA pre-alignment references were retrieved and built using Refgenie^80^. Samples were processed with adapter trimming, alignment with Bowtie2^81^, removal of rDNA, quality control, and generation of strand-specific signal tracks. Output bigwig files were prepared at 1bp resolution with combined positive and negative strands. deeptools^82^v3.5.1 was used to display metagene analysis.

### CUT&RUN

CUT&RUN was performed using the EpiCypher CUTANA CUT&RUN Kit v1.0 per the manufactures protocol with no deviations. Approximately 500k cells/condition were used, pretreated with 1 μM GSK591 for 0-, 4-, or 7-days. IgG negative controls were performed on DMSO-treated cells at 0 or 7 days. The following antibodies were used: IgG Negative Control (Epicypher 13-0042); H3K4me3 (Epicypher 13-0060); H3K27me3 (Thermo Fisher MA5-11198). Library prep was performed with the CUTANA CUT&RUN Library prep kit according to the manufacturer’s instructions. Fragment size was confirmed using Bioanlyzer Analysis and then sequenced on an Illumina NextSeq 500 with paired reads of read length 35. Reads were mapped to hg38 using bowtie2^81^, peaks called using macs2 and output bigwig files normalized using counts per million (CPM) mapped reads, all in the nfcore/cutandrun v3.2.1 pipeline. For publicly available ChIP-Seq data, reads were mapped to hg38 using bwa and peaks called using macs3 in the nf-core/chipseq v2.1.0 pipeline. Visualization was performed in Integrative Genomics Viewer (IGV) v2.14.1.

### Lentiviral CRISPRi knockdowns

Protocol was completed as reported previously^53^. The lenti_dCas9-KRAB-MeCP2 plasmid was a gift from Andrea Califano (https://www.addgene/122205; RRID: Addgene_122205) and pXPR_050 was a gift from John Doench & David Root (https://www.addgene/96925; RRID: Addgene_96925)^83, 84^. Custom sgRNA sequences (**Supplementary Table S5**) were designed using the Broad CRISPick software^85,86^(https://portals.broadinstitute.org/gppx/crispick/public) and then cloned into pXPR_050 as previously described^84^.

Lentiviral particles were generated and transduced into cells by spinfection as previously reported^35^. Stable expression of the dCas9-KRAB-MeCP2 construct was achieved in A549 cells with Blasticidin (Cayman) selection at 10 μg/mL. sgRNA were transduced by spinfection, grown for 24-hours, then selected with 2 μg/mL puromycin (Cayman). Cells were harvested on day 7 post-spinfection for downstream analyses.

K562 cells were nucleofected with the dCas9-KRAB-MeCP2 construct and stable lines were generated with 500 μg/mL blasticidin (Cayman). For each experiment, cells were nucleofected with sgRNA lentiviral particles, then maintained in standard growth media for 48 hours, followed by selection with 2 μg/mL puromycin. Cells were harvested on day 5 post-nucleofection for downstream analyses.

### shRNA-mediated prmt5 knockdown

Mouse embryonic stem cells (mESCs) were subjected to shRNA-mediated knockdown of Prmt5 using a Lipofectamine 3000-based reverse transfection protocol. On day 0, cells were transfected with shRNA-encoding plasmids (gift from Dr. Andrew Hutchins, Southern University of Science and Technology, China^87^) complexed with Lipofectamine 3000 in antibiotic-free media. Cells were cultured for 48 hours post-transfection without selection. From day 2 to day 3, cells were treated with 2 μg/mL puromycin to enrich for transfected populations. Cells were harvested on day 4 for downstream analyses.

### Cellular senescence assay

Cells were lentivirally transduced for p53 knockdown, selected for 3 days in puromycin, then plated in 6-well plates and treated with 1μM GSK591 or (-)-Nutlin-3 for 4-days or 0.5μM Doxorubicin for 24-hours followed by 3-day washout. β-galactosidase senescence staining was performed according to the manufacturer’s protocol (Cell Signaling Technology; 9860S).

### Palbociclib G1 arrest

A549 cells were plated in assay plates containing 0.5μM palbociclib and grown for 30 hours (duration of one A549 cell cycle) to arrest in G1 phase, then released into S phase following one rinse with warmed PBS, and addition of media ±1μM GSK591. At the appropriate time, cells were removed to ice, rinsed with PBS, and processed with the Monarch Total RNA kit (New England Biolabs) and as described above for general RT-qPCR.

### Immunoblotting for soluble histones

Soluble histone immunoblots were completed as immunoblots above with altered extraction buffer and no sonication: 1% TritonX-100, 0.1% SDS, 150mM NaCl, 1mM EDTA, 50mM pH 7.8 Tris-HCl, protease and phosphatase inhibitors.

### Subcellar fractionation and proteomics using Tan-dem-Mass-Tag labeling

A549 cells were harvested by trypsinization, washed in 37°C PBS, and resuspended at a concentration of 1.5×10^6^ cells/mL in complete DMEM. Cells were plated at appropriate densities to account for cell proliferation and treated with 1 μM GSK591, 1 μM MS023, or 0.01% DMSO for 15 min, 90 min, 180 min, or 48 hours. At each time point, cells were washed with 37 °C PBS containing inhibitor and trypsinized followed by centrifugation at 300g for 3 min at 4°C, resuspended in 1 mL cold PBS, and transferred to low-retention, low-adhesion tubes. Cell pellets were resuspended in 300 μL hypotonic lysis buffer (10 mM Tris-HCl pH 8.0 at 4°C, 0.1% NP-40, 1 mM KCl, 1.5 mM MgCl_2_, protease and phosphatase inhibitors, 40 U/mL RNaseOUT, and 1 μM PRMT inhibitor) using a wide-orifice pipette tip. The suspension was rotated end-over-end for 30 min at 4°C. Lysates were then centrifuged at 10,000 × g for 10 min at 4°C to separate cytosolic and nuclear fractions. The supernatant containing the cytosolic fraction was then isolated and an aliquot taken for western blot analysis. Nuclear pellets were washed once with hypotonic buffer and resuspended in 300 μL nuclear lysis buffer (10 mM Tris-HCl pH 8.0 at 4°C, 0.1% NP-40, 400 mM KCl, 1 mM DTT, protease and phosphatase inhibitors, 40 U/mL RNaseOUT, and 1 μM PRMT inhibitor). The suspension was rotated for 30 min at 4°C and centrifuged at 10,000g for 10 min. The supernatant, containing nucleoplasmic proteins, was processed analogously to the cytoplasmic fraction. The chromatin pellet was washed once with nuclear lysis buffer and then resuspended in 150 μL nuclear lysis buffer and subjected to probe-tip sonication at 20% amplitude for 5 seconds. Protein concentrations were determined using the BCA assay and diluted to a final 100 μg per 100 μL followed by nucleic acid digestion with Benzonase (Millipore Sigma; 70664).

### S-trap protein digestion for TMT fractionation

Fractionated extracts were amended to 5% SDS, 5 mM DTT and 50 mM ammonium bicarbonate (pH = 8), and left on the bench for about 1 hour for disulfide bond reduction. Samples were then alkylated with 20 mM iodoacetamide in the dark for 30 minutes. Afterward, phosphoric acid was added to the sample at a final concentration of 1.2%. Samples were diluted in six volumes of binding buffer (90% methanol and 10 mM ammonium bicarbonate, pH 8.0). After gentle mixing, the protein solution was loaded to an S-trap filter (Protifi) and spun at 500 g for 30 sec. The sample was washed twice with binding buffer. Finally, 1 μg of sequencing grade trypsin (Promega), diluted in 50 mM ammonium bicarbonate, was added into the S-trap filter and samples were digested at 37°C for 18 hours. Peptides were eluted in three steps: (i) 40 μl of 50 mM ammonium bicarbonate, (ii) 40 μl of 0.1% TFA and (iii) 40 μl of 60% acetonitrile and 0.1% TFA. The peptide solution was pooled, spun at 1,000 g for 30 sec and dried in a vacuum centrifuge.

For TMT labeling, samples were resuspended in 100 mM triethylammonium bicarbonate (TEAB, pH 8.5) and labeled with TMT reagents (Thermo Scientific: 90111) according to the manufacturer’s instructions. After a 1-hour incubation at room temperature, the reaction was quenched with 5% hydroxylamine for 15 minutes. Labeled peptides were pooled, dried, and subjected to C18 desalting prior to LC-MS/MS analysis.

### Palbociclib G1 arrest and release for SILAC

A549 cells were grown in DMEM for SILAC (ThermoFisher: 88364) supplemented with light L-Lysine (ThermoFisher: 89987), L-Arginine HCl (ThermoFisher: 89989), and 10% charcoal-stripped FBS (ThermoFisher: A3382101) for one week, then plated in assay plates containing 0.5μM palbociclib and grown for 30 hours (duration of one A549 cell cycle) to arrest in G1 phase. To release into S phase, cells were rinsed with warm PBS, then incubated in media +/-1μM GSK591 containing heavy L-Lysine (^13^C_6_-^15^N_2_) (ThermoFisher: 88209) in place of of light L-Lysine. After 9 hours, cells were rinsed with chilled PBS and harvested by scraping, collected by centrifugation, and pellets frozen at −80°C for subsequent processing.

### Total proteome sample preparation for SILAC

Total proteome cell pellets were resuspended with 50ul of 1% SDS in 50mM ammonium bicarbonate pH 8.0 containing 1ul/reaction Pierce universal nuclease (ThermoFisher: 88701), followed by addition of 5mM DTT (dithiothreitol, Sigma) and one hour incubation at 37ºC. Finally, iodoacetamide (Sigma) was added to 20mM and incubated for 30 minutes at RT. Prior to loading onto Strap filters (Protifi), phosphoric acid was added to a concentration of 1.2% and samples were diluted in Strap binding buffer (90% methanol and 10mM ammonium bicarbonate). Samples on S-trap filters were washed three times with binding buffer then treated with 1.1ug of sequencing grade LysC (NEB) pre-pared in 50mM ammonium bicarbonate pH 8.0 and digested overnight at 37ºC. Digested peptides were collected with sequential elutions with 50mM ammonium bicarbonate, 0.1% trifluoroacetic acid (TFA), and 0.1% TFA/60% acetonitrile, then dried by speed vacuum.

### Sample desalting for all mass spectrometry samples

Prior to mass spectrometry analysis, samples were desalted using a 96-well plate filter (Orochem) packed with 1 mg of Oasis HLB C-18 resin (Waters). Briefly, the samples were resuspended in 100 μl of 0.1% TFA and loaded onto the HLB resin, which was previously equilibrated using 100 μl of the same buffer. After washing with 100 μl of 0.1% TFA, the samples were eluted with a buffer containing 70 μl of 60% acetonitrile and 0.1% TFA. Peptides were dried by speed vacuum and stored at −80ºC until analysis.

### LC-MS/MS acquisition and analysis

Samples were resuspended in 10 μl of 0.1% TFA and loaded onto a Dionex RSLC Ultimate 300 (Thermo Scientific), coupled online with an Orbitrap Fusion Lumos (Thermo Scientific). Chromatographic separation was performed with a two-column system, consisting of a C-18 trap cartridge (300 μm ID, 5 mm length) and a picofrit analytical column (75 μm ID, 25 cm length) packed inhouse with reversed-phase Repro-Sil Pur C18-AQ 3 μm resin. Peptides were separated using a 90 min gradient from 4-30% buffer B (buffer A: 0.1% formic acid, buffer B: 80% acetonitrile + 0.1% formic acid) at a flow rate of 300 nl/min. The mass spectrometer was set to acquire spectra in a data-dependent acquisition (DDA) mode. Briefly, the full MS scan was set to 300-1200 m/z in the orbitrap with a resolution of 120,000 (at 200 m/z) and an AGC target of 5×10^5^. MS/MS was performed in the ion trap for label-free experiments (SILAC) and in the orbitrap for the TMT experiment.

Proteome raw files were searched using Proteome Discoverer software (v2.5, Thermo Scientific) using SEQUEST search engine and the SwissProt human database. The search for total proteome included variable modification of N-terminal acetylation, and fixed modification of carbamidomethyl cysteine. Trypsin or LysC was specified as the digestive enzyme with up to 2 missed cleavages allowed. Mass tolerance was set to 10 pm for precursor ions and 0.2 Da for product ions. Peptide and protein false discovery rate was set to 1%. Following the search, data was processed as described previously^88^. Briefly, proteins were log_2_ transformed, normalized by the average value of each sample, and missing values were imputed using a normal distribution 2 standard deviations lower than the mean. For SILAC samples, the heavy/light ratio was calculated without normalization. Statistical regulation was assessed using heteroscedastic t-test (if p-value < 0.05). Data distribution was assumed to be normal, but this was not formally tested.

### Analysis of histones by high-performance liquid chromatography

Histones were acid-extracted as reported in the previous section. Samples were filtered through a 0.2 μm Millex syringe filter into HPLC vials and analyzed using a Waters 2695 Separations Module with a Vydac 218TP C18 column. Mouse samples were run using the method set methodMAW_Scout_Col250×1_21_long93 (93 min run time). Human samples included an additional two methanol washes prior to injection. Eluted peaks were monitored at 214 nm using a Waters 996 Photodiode Array Detector. H1 peak areas were quantified using Waters Empower Pro software (v.2) and normalized to the corresponding H2B peak areas.

### Expression of methylarginine mutants

Histone H4 constructs (H4C14) were purchased from GenScript, PCR amplified to insert a C-terminal HA tag and individually cloned into a pDONR221 vector of the Gateway lentiviral system using the manufacture’s protocols (Invitrogen). Entry clones were cloned into pLEX_307 (a gift from David Root; https://www.addgene.org/41392; RRID: Addgene_41392) with premature stop codon to exclude the v5 tag. Sequences are found in **Supplemental Table 5**. Lentiviral particles were produced as reported previously and transduced cells were selected in 2 μg/ml puromycin (Cayman).

### Statistical analysis

All immunoblots were performed in at least two independent biological replicates. Statistical analyses were performed either using GraphPad Prism (v10.1.0) or R (v4.4.0). Independent t-tests were performed to compare means between only two groups.

## Supporting information

Supplemental Table S1 - PROSeq DESeq2

Supplemental Table S2 - TFEA PROSeq Analysis

Supplemental Table S3 - TMT Fractionated Proteomics

Supplemental Table S4 - SILAC S-phase Proteomics

## Data Availability

Raw data for sequencing experiments were deposited under GEO: PRO-seq (GSE275217 and GSE275220), CUT&RUN (pending accession number). Raw data for total cell and fractionated LC-MS/MS is deposited to the ProteomeXchange Consortium via the PRIDE partner repository: PXD065294. All code and FIJI macros used to generate data in this manuscript can be found here: https://github.com/Shechterlab/Roth_etal_2025.

## Acknowledgments

The histone H3 and H4 antibodies were a generous gift from C. David Allis. This work was supported by the National Institutes of Health [T32GM149364 to J.S.R., D.L.Y. and J.D.D., R01GM108646 to D.S., R01GM147165 to A.S.], Irma T. Hirschl and Monique Weill-Caulier Charitable Trusts to D.S., and The ALS Therapy Development Institute to D.S. The Sidoli lab gratefully acknowledges for funding the Hevolution Foundation (AFAR), the Einstein-Mount Sinai Diabetes center, and the NIH Office of the Director (S10OD030286). We thank Emmanuel Burgos for early studies. JSR would like to thank Matt Wooten for exciting and valuable discussions.

## Author contributions

J.S.R. and D.S. conceived of the study. A.I.S. conceived of experiments. J.S.R., J.D.D., D.Y., M.I.M., A.S, H. P., S.H., J.B., an N.J completed experiments. J.S.R., M.I.M., J.T.A, V.G., C.C.Q, J.B., J.A., S.S., and D.S. performed data analysis. J.S.R. and D.S. wrote the manuscript. All authors approved the manuscript. To improve concision in portions of this manuscript, during the preparation of this work the authors used ChatGPT4.0o and ChatGPT4.5. After using this tool, the authors fully reviewed and edited the content as needed and take full responsibility for the content of the publication.

