## Supplemental Figures S1-S7 for "PRMT5 activity sustains histone production to maintain genome integrity"

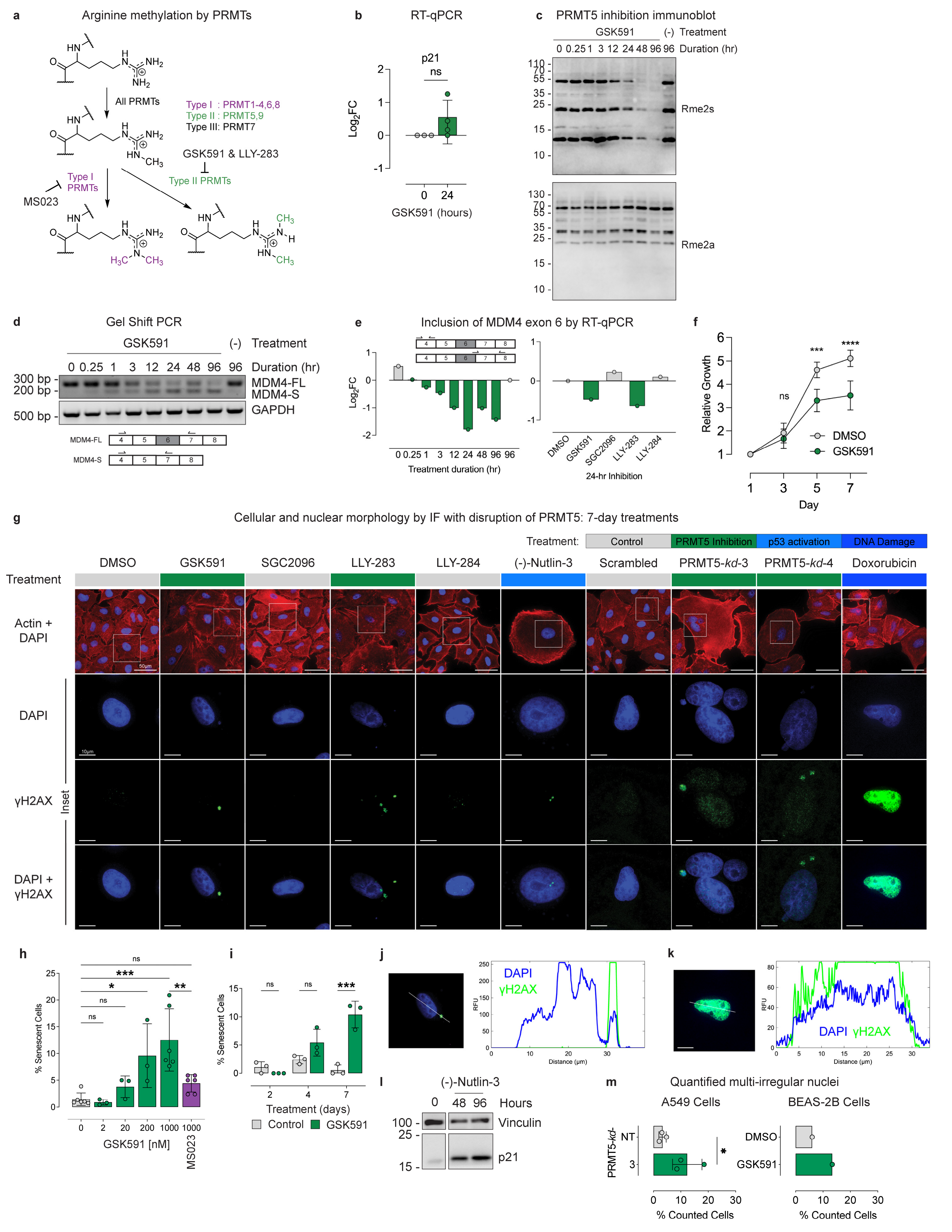
a
Arginine methylation by PRMTs
All PRMTs
Type I : PRMT1-4,6,8
Type II : PRMT5,9
Type III: PRMT7
GSK591 & LLY-283
Type II PRMTs
Type I PRMTs
MS023
b
RT-qPCR
p21
ns
Log2FC
GSK591 (hours)
c
PRMT5 inhibition immunoblot
GSK591
(-) Treatment
0 0.25 1 3 12 24 48 96 96 Duration (hr)
Rme2s
Rme2a
d
Gel Shift PCR
GSK591
(-) Treatment
0 0.25 1 3 12 24 48 96 96 Duration (hr)
300 bp
200 bp
500 bp
MDM4-FL
MDM4-S
GAPDH
e
Inclusion of MDM4 exon 6 by RT-qPCR
Log2FC
Treatment duration (hr)
DMSO
GSK591
SGC2096
LLY-283
LLY-284
24-hr Inhibition
f
Relative Growth
ns
***
****
DMSO
GSK591
Day
g
Cellular and nuclear morphology by IF with disruption of PRMT5: 7-day treatments
Treatment:
Control
PRMT5 Inhibition
p53 activation
DNA Damage
DMSO
GSK591
SGC2096
LLY-283
LLY-284
(-)-Nutlin-3
Scrambled
PRMT5-kd-3
PRMT5-kd-4
Doxorubicin
Treatment
Actin + DAPI
DAPI
γH2AX
Inset
DAPI + γH2AX
50µm
10µm
h
% Senescent Cells
ns
***
*
**
GSK591 [nM]
MS023
i
% Senescent Cells
ns
***
Treatment (days)
Control
GSK591
j
DAPI
γH2AX
RFU
Distance (µm)
k
DAPI γH2AX
RFU
Distance (µm)
l
(-)-Nutlin-3
0 48 96 Hours
Vinculin
p21
m
Quantified multi-irregular nuclei
A549 Cells
BEAS-2B Cells
PRMT5-kd-
NT
DMSO
GSK591
% Counted Cells

**Supplemental Figure S1 (Related to Figure 1). PRMT5 inhibition rapidly induces alternative splicing. a)** Schematic of arginine methylation by the family of PRMTs. GSK591 and LLY-283 inhibit PRMT5; MS023 inhibits type I PRMTs. **b)** RT-qPCR of p21 induction after 24-hours GSK591 in A549 cells. **c)** Immunoblots of total cellular lysate after PRMT5 inhibition. Rme2s blot from figure 1a is cropped from S1c. **d)** Gel shift PCR tracking alternative splicing of MDM4 over time course of GSK591 treatment. **e)** Quantitative RT-qPCR tracking alternative splicing of MDM4 over time course of GSK591 treatment and 24-hours of GSK591 and LLY-283 but not structurally similar negative controls SGC2096 or LLY-284. **f)** Growth curve measuring metabolic activity of A549 cells treated with GSK591; n=4 biological replicates in A549 cells. **g)** Representative images displaying cellular phenotypic changes and micronuclei formation upon PRMT5 activity disruption, with relevant negative controls. For the sake of comparison, IF images from Figure 1 for DMSO, GSK591, and LLY-283 are repeated here. Scale bar 50μm in Actin+DAPI and 10μm in inset images. **h)** Dose response of GSK591 treatment on β-galactosidase senescence in A549 cells. **i)** Time course of GSK591 treatment on β-galactosidase senescence in A549 cells. **j)** Profile plot of DAPI (blue) and γH2AX (green) intensity across a representative nucleus after 7-days of PRMT5 inhibition with GSK591, highlighting a γH2AX-positive micronuclei. **k)** Representative image of A549 cells treated with 24-hours of 1uM doxorubicin, highlighting pan-nuclear γH2AX (green) staining within nuclei (blue) with profile plot of DAPI (blue) and γH2AX (green) intensity across a representative nucleus. **l)** Quantified multi-irregular nuclei in A549 cells after 7-day PRMT5 knockdown; n=3 biological replicates with >100 cells counted/replicate, and in non-cancerous BEAS-2B lung cells after 4-days of GSK591 treatment; n=1 with >100 cells counted. **m)** Immunoblot validation of p21 induction by p53 activation via (-)-Nutlin-3 treatment in A549 cells. Student's one-tailed t-tests were applied; p-value = *<0.05, **<0.005, ***<0.0005, ****<0.00005, n.s. = not significant.

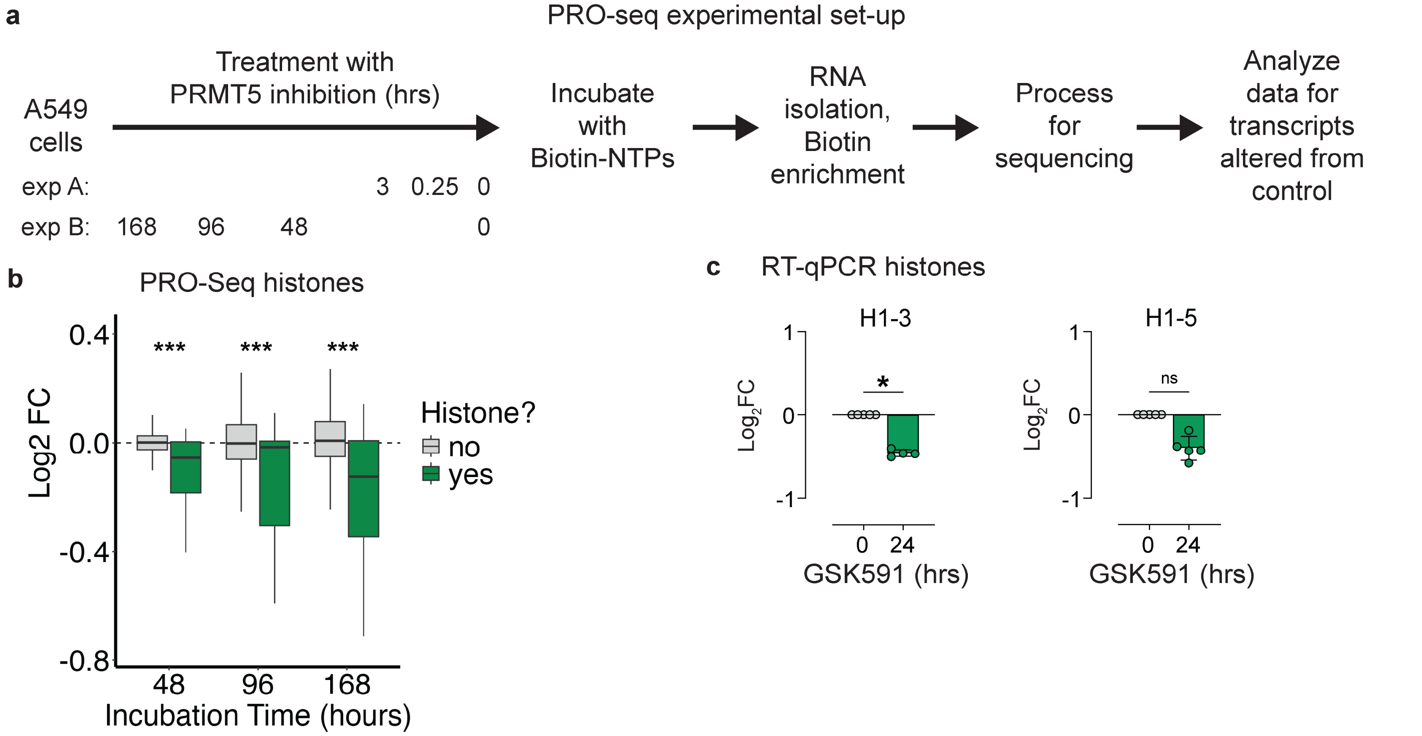


**Supplemental Figure S2 (Related to Figure 2). PRO-seq experimental set-up. a)** Experimental set-up for PRO-Seq studies. **b)** Random permutation testing of $Log_2FC$ nascent transcription response for all histone genes vs all other genes at 48-, 96-, and 168-hours of PRMT5 inhibition (100,000 permutations). p-value = ***<0.0005. **c)** RT-qPCR quantification of two representative histone genes after 24-hours of PRMT5 inhibition in asynchronous cells. Student's one-tailed t-test was applied. p-value = *<0.05, n.s. = not significant.

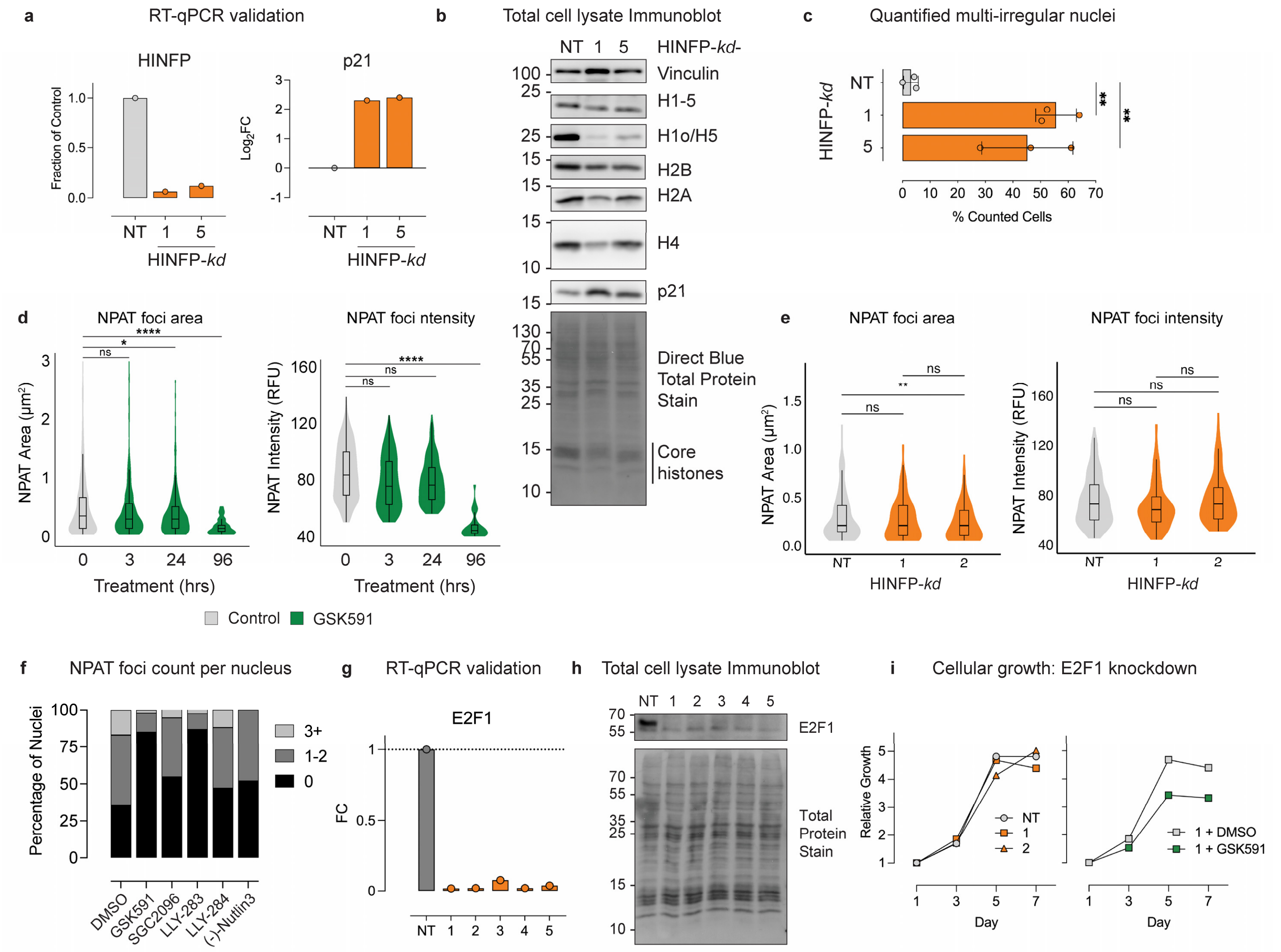


**Supplemental Figure S3 (Related to Figure 3). HINFP knockdown validation**. **a)** RT-qPCR validation of 4-day HINFP knockdown and resultant p21 induction. **b)** Immunoblots of total cellular lysate 4-days after HINFP knockdown. **c)** Quantification of multi-irregular nuclei after 7-days of HINFP knockdown in A549 cells; n=3 biological replicates. **d)** NPAT foci area and intensity quantified from 4-day PRMT5 inhibition in A549 cells; n=1 biological replicate with >200 nuclei counted/condition for GSK591 time course. **e)** NPAT foci area and intensity quantified from 4-day HINFP knockdown in A549 cells; n=2 biological replicates with >75 nuclei counted/condition for HINFP knockdown. **f)** Count of NPAT foci per nucleus from treatments in figure 1f; n=3 biological replicates with >100 nuclei counted/replicate. **g)** RT-qPCR validation of 7-day E2F1 knockdown. **h)** Total cell lysate immunoblot validation 7-days after E2F1 knockdown. **i)** Growth curve measuring metabolic activity of A549 cells after E2F1 knockdown; n=1 biological replicate. Student's one-tailed t-tests were applied; p-value = *<0.05, **<0.005, ***<0.0005, ****<0.00005, n.s. = not significant.

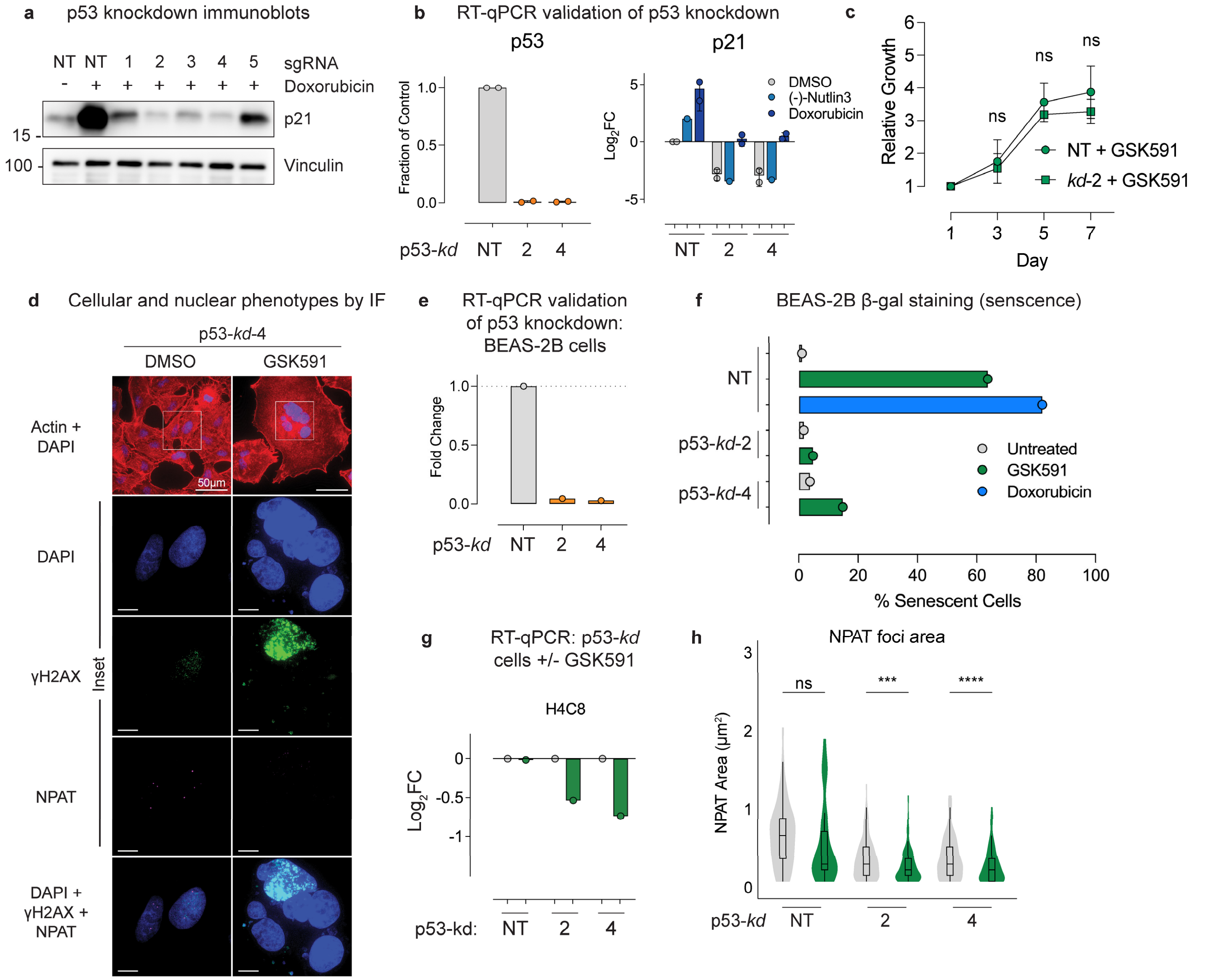


**Supplemental Figure S4 (Related to Figure 4). p53 knockdown validation. a-b)** Immunoblot and RT-qPCR validation after 4-day p53 knockdown in A549 cells ± 24-hours 1µM doxorubicin or (-)-Nutlin-3. **c)** Growth curve directly comparing PRMT5 inhibition between non-targeting (NT) sgRNA and p53 knockdown; mean ±SD, n=2 biological replicates. **d)** Immunofluorescence staining of p53-*kd*-4 ± 4-day PRMT5 inhibition. **e)** RT-qPCR validation after 4-day p53 knockdown in BEAS-2B cells. **f)** β-galactosidase staining positivity of p53 knockdown cells ± PRMT5 inhibition in BEAS-2B cells. **g)** RT-qPCR analysis of H4C8 in p53 knockdown A549 cells treated with 24-hours PRMT5 inhibition. **h)** NPAT foci area quantifications from p53 knockdown A549 cells from figures 4f-g. Student's one-tailed t-tests were applied; p-value = ***<0.0005, ****<0.00005, n.s. = not significant.

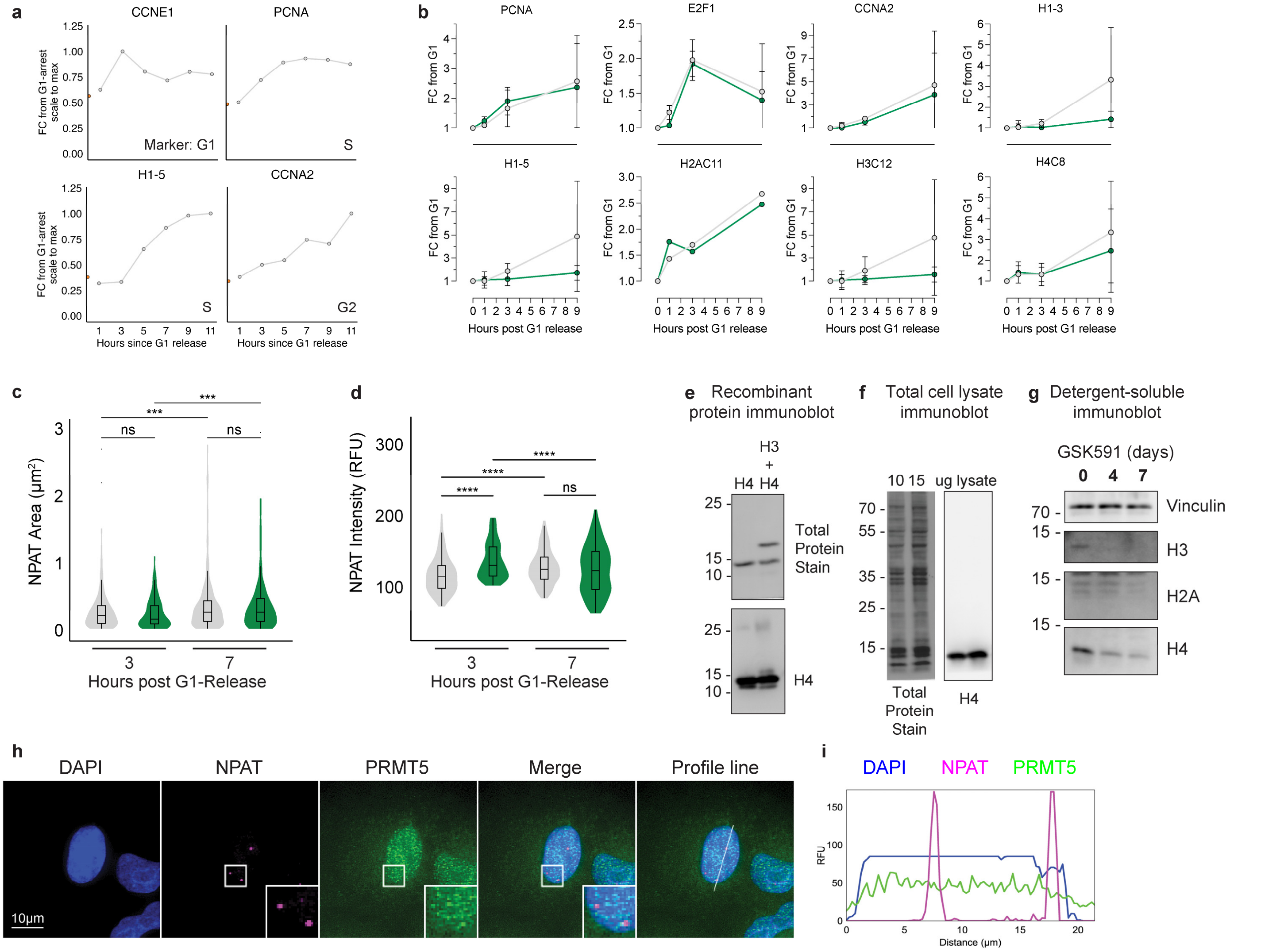


**Supplemental Figure S5 (Related to Figure 5). S-phase progression validation and controls. a)** S phase progression validation after Palbociclib arrest and release; data represent fold-changes from palbociclib release, scaled to maximum value over the time-course; n=2 biological replicates. **b)** Fold change response of various cell cycle markers and histones over 9 hours of release into S phase ± PRMT5 inhibition. **c-d)** NPAT foci area and intensity throughout S phase release ± PRMT5 inhibition; n=1 biological replicate, >200 cells counted per condition. Student's one-tailed t-tests were applied; p-value = ***<0.0005, ****<0.00005, n.s. = not significant. **e)** Validation of H4 antibody (Gifted by C. D. Allis) on recombinant protein. **f)** Validation of H4 antibody on A549 cellular lysate. **g)** Immunoblot for soluble histones after PRMT5 inhibition. **h)** Immunofluorescence staining of NPAT and PRMT5 in A549 cells. **i)** Profile line plots of representative NPAT foci (HLBs) and PRMT5 intensity.

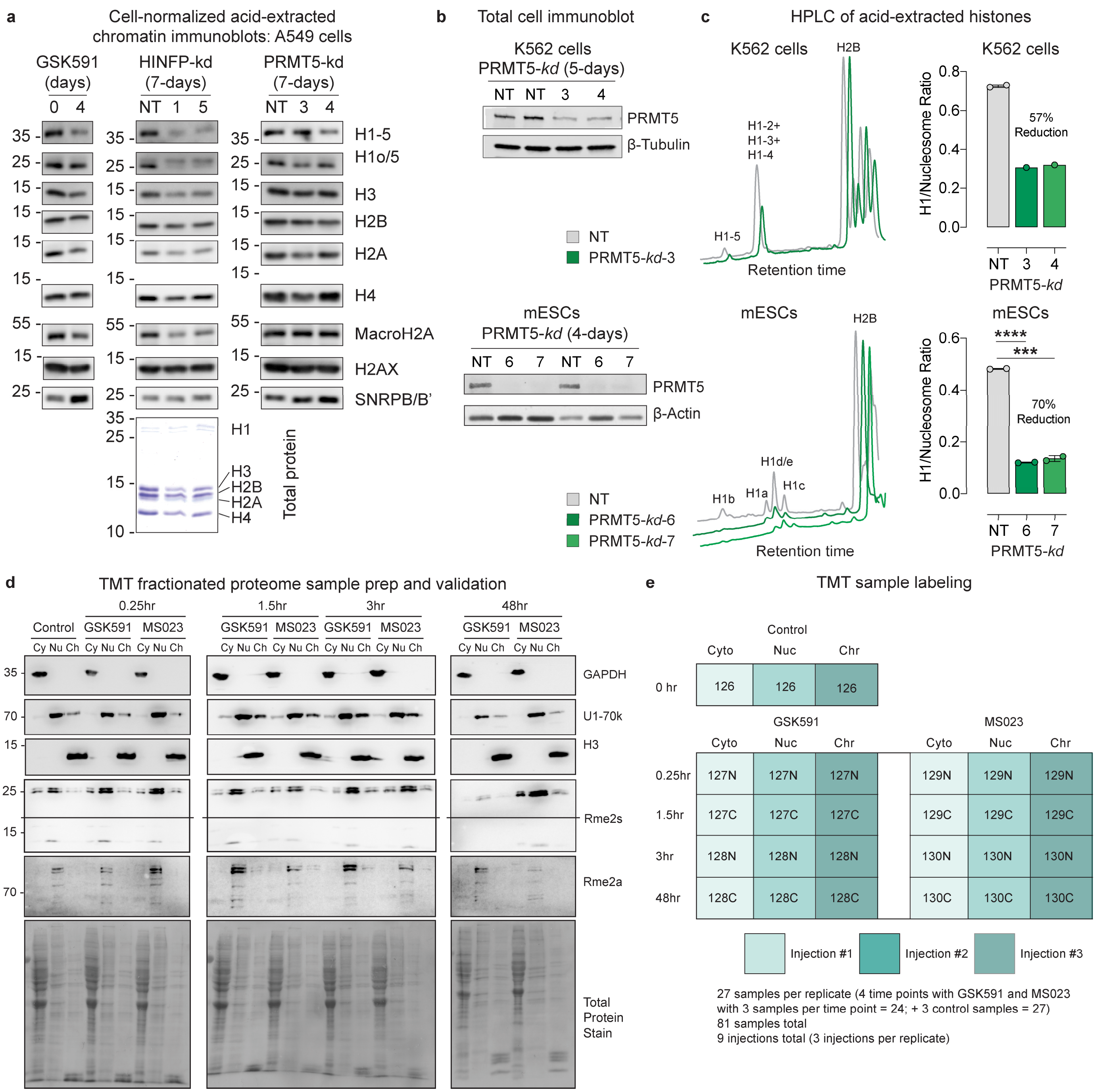


**Supplemental Figure S6 (Related to Figure 6). TMT labeling mass spectrometry controls. a)** Immunoblots of acid-extracted chromatin from equal cell number in A549 cells; left: 4 days of PRMT5 inhibition, middle: 7 days HINFP knockdown, right: 7 days PRMT5 knockdown. **b)** Immunoblot confirming PRMT5 knockdown in whole-cell lysates of (top) K562 cells and (bottom) mESCs. **c)** Left: HPLC chromatogram of acid-extracted histones (equal protein input) from non-targeting control and PRMT5 knockdown cells showing a reduction in the H1/nucleosome ratio. Top: K562 cells; Bottom: mESCs. Right: Quantification of H1/nucleosome peak area following PRMT5 knockdown. Top: K562 cells; Bottom: mESCs. Student's one-tailed t-tests were applied; p-value = ***<0.0005, ****<0.00005, n.s. = not significant. **d)** Cell fractionation validation by immunoblotting for TMT labeling study. **e)** TMT sample labeling set-up.

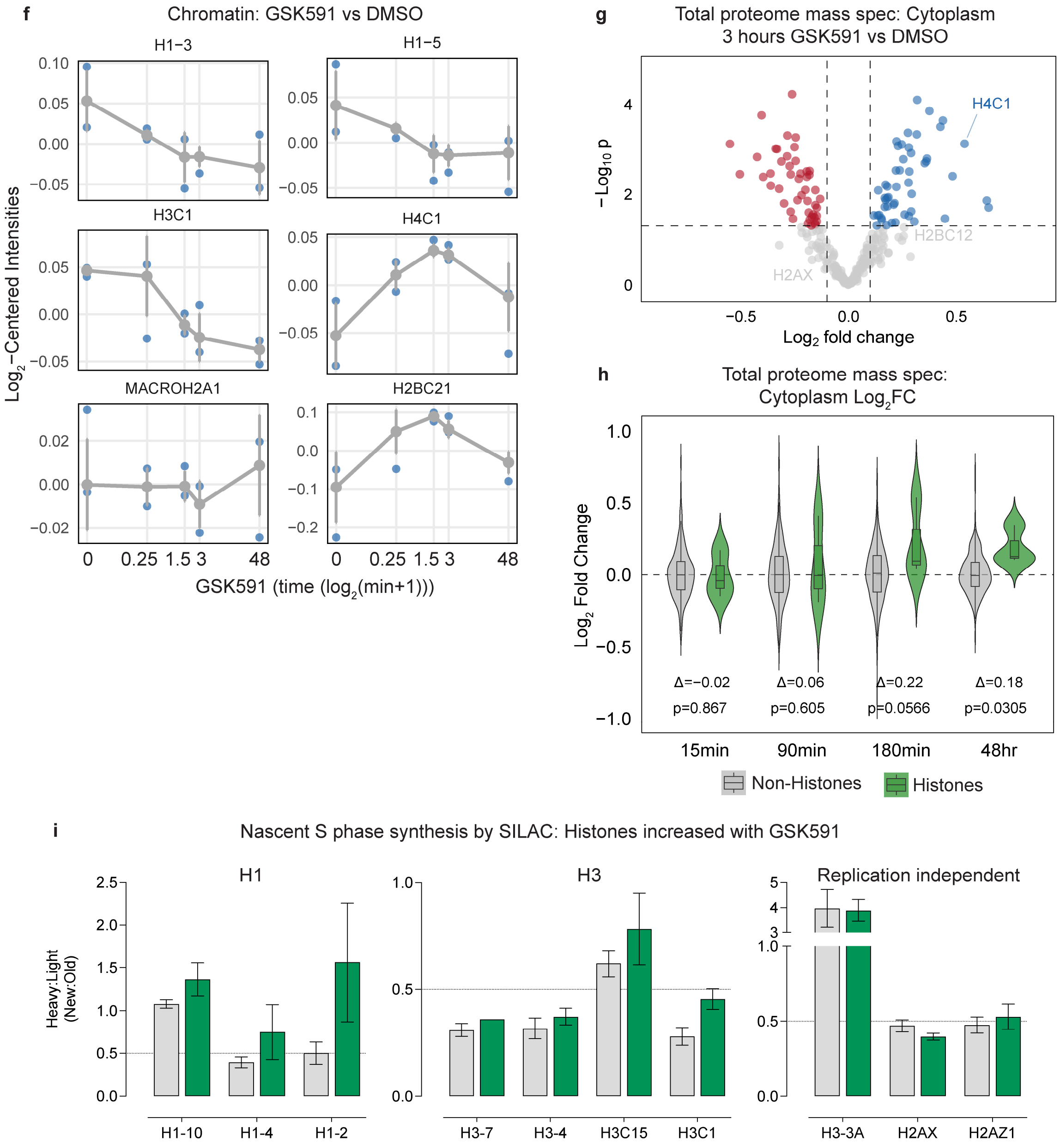


**Supplemental Figure S6 continued (Related to Figure 6). SILAC data continued. f)** Quantification of selected histones from the chromatin fraction over the PRMT5 inhibition time-course; $Log_2$-centered intensity from 4 replicates at each timepoint. **g)** Total proteome mass spectrometry on the cytoplasmic fraction. **h)** Non-histone vs histone $Log_2FC$ over time; cytoplasmic fraction. Random permutation test (10,000). **i)** Quantified nascent histones by SILAC after 9-hour release into S phase. Bars represent mean ±SEM of 4 replicates.

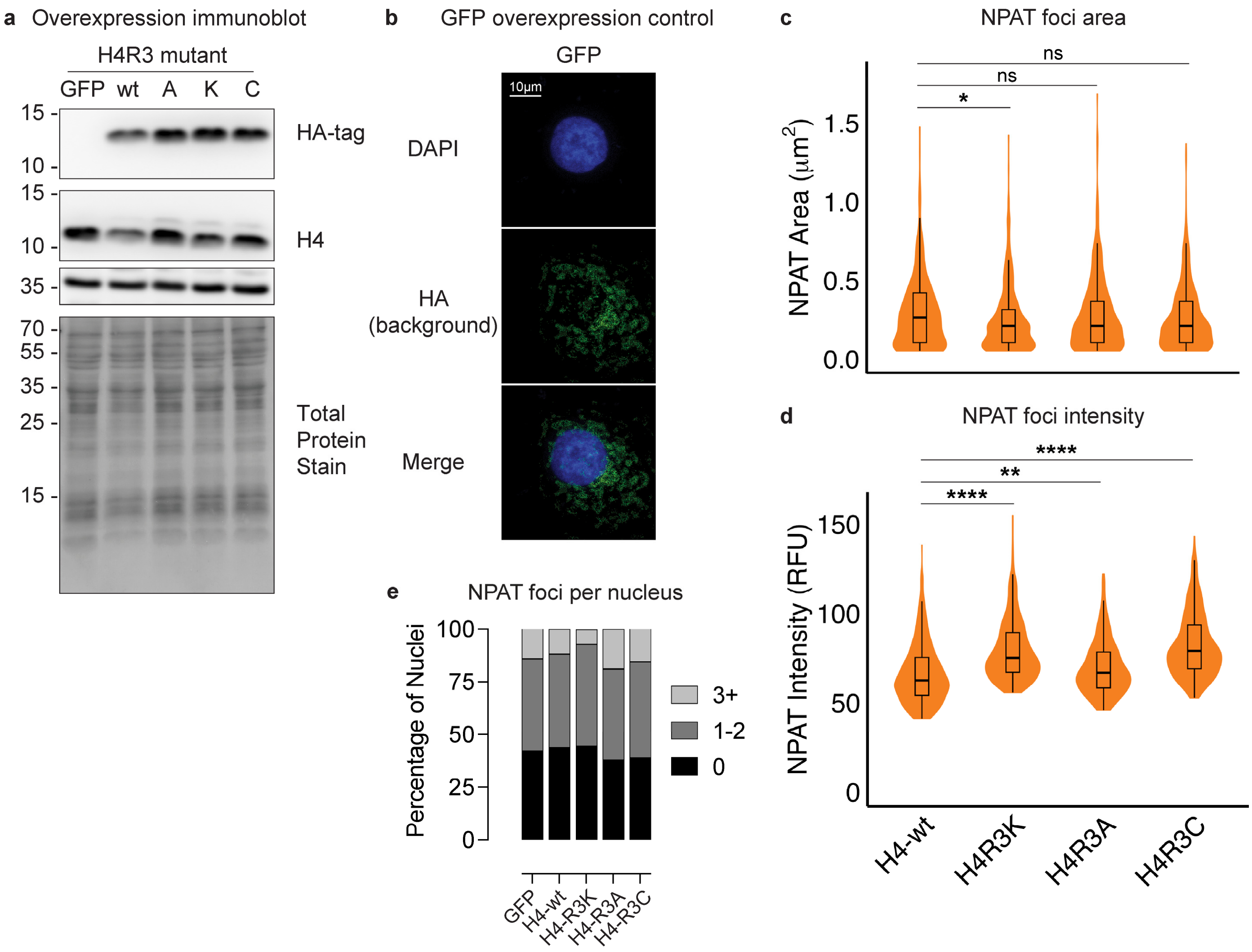


**Supplemental Figure S7 (Related to Figure 7). Mutant histone overexpression validation. a)** Immunoblot validation of HA-tagged histone H4 overexpression in A549 cells. **b)** IF for background HA signal in A549 cells overexpressing GFP without HA tag. **c-e)** NPAT foci area, intensity, and count from mutant histone H4 overexpression; n=1 biological replicates, >225 cells/condition. Student's one-tailed t-tests were applied; p-value = *<0.05, **<0.005, ***<0.0005, ****<0.00005, n.s. = not significant.
